# Global ubiquitylation analysis of mitochondria in primary neurons identifies physiological Parkin targets following activation of PINK1

**DOI:** 10.1101/2021.04.01.438131

**Authors:** Odetta Antico, Alban Ordureau, Michael Stevens, Francois Singh, Marek Gierlinski, Erica Barini, Mollie L. Rickwood, Alan Prescott, Rachel Toth, Ian G. Ganley, J. Wade Harper, Miratul M. K. Muqit

## Abstract

Autosomal recessive mutations in PINK1 and Parkin cause Parkinson’s disease. How activation of PINK1 and Parkin leads to elimination of damaged mitochondria by mitophagy is largely based on cell culture studies with few molecular studies in neurons. Herein we have undertaken a global proteomic-analysis of mitochondria from mouse neurons to identify ubiquitylated substrates of endogenous Parkin activation. Comparative analysis with human iNeuron datasets revealed a subset of 49 PINK1-dependent diGLY sites upregulated upon mitochondrial depolarisation in 22 proteins conserved across mouse and human systems. These proteins were exclusively localised at the mitochondrial outer membrane (MOM) including, CISD1, CPT1α, ACSL1, and FAM213A. We demonstrate that these proteins can be directly ubiquitylated by Parkin *in vitro*. We also provide evidence for a subset of cytoplasmic proteins recruited to mitochondria that undergo PINK1 and Parkin independent ubiquitylation including SNX3, CAMK2α and CAMK2β indicating the presence of alternate ubiquitin E3 ligase pathways that are activated by mitochondrial depolarisation in neurons. Finally we have developed an online resource to visualise mitochondrial ubiquitin sites in neurons and search for ubiquitin components recruited to mitochondria upon mitochondrial depolarisation, MitoNUb. This analysis will aid in future studies to understand Parkin activation in neuronal subtypes. Our findings also suggest that monitoring ubiquitylation status of the 22 identified MOM proteins may represent robust biomarkers for PINK1 and Parkin activity *in vivo*.

## INTRODUCTION

Mitochondria perform diverse functions within eukaryotic cells that are essential to their survival however under physiological conditions, they are exposed to pleiotropic stress including reactive oxygen species (ROS) and misfolded and aggregated proteins that cause mitochondrial dysfunction (Vafai and Mootha, 2012, Youle, 2019). This is particularly relevant in terminally differentiated cell types such as neurons in which aberrant mitochondrial homeostasis is linked to diverse neurological syndromes and neurodegenerative disorders (McFarland et al., 2010, Johri and Beal, 2012). Mitochondrial quality control pathways have evolved to sense and protect against mitotoxic stress, of which elimination to lysosomes via the autophagy pathway (mitophagy) has been extensively studied in recent years (Montava-Garriga and Ganley, 2020, Youle, 2019). Autosomal recessive mutations in human PTEN-induced kinase 1 (PINK1) and the RING-IBR-RING (RBR) ubiquitin E3 ligase Parkin (encoded by *PARK6* and *PARK2* genes respectively) are causal for early-onset Parkinson’s disease (PD) (Kitada et al., 1998, Valente et al., 2004). Landmark cell-based studies have demonstrated that these proteins function within a common pathway to regulate stress-evoked mitophagy via a ubiquitin-dependent mechanism (Harper et al., 2018, Singh and Muqit, 2020, Moehlman and Youle, 2020). Upon loss of mitochondrial membrane potential that can be induced artificially by mitochondrial uncouplers (e.g. Antimycin A/Oligomycin (AO)), PINK1 becomes stabilised and activated on the mitochondrial outer membrane (MOM) where it phosphorylates both ubiquitin and Parkin at their respective Serine65 (Ser65) residues leading to stepwise recruitment and activation of Parkin at the OMM (Kane et al., 2014, Kazlauskaite et al., 2014b, Kondapalli et al., 2012, Koyano et al., 2014, Ordureau et al., 2014). An analogous pathway is activated upon accumulation of misfolded proteins in the mitochondrial matrix or upon mitochondrial damage occurring in cells lacking Mitofusin proteins (MFN1 and MFN2), indicative of physiologically relevant mitochondrial stress pathways (Burman et al., 2017, Narendra et al., 2008). Active Parkin ubiquitylates myriad substrates including Voltage-dependent anion channels (VDACs), MitoNeet/CDGSH iron-sulfur domain-containing protein 1 (CISD1), Mitofusins (MFNs), Hexokinase 1 (HK1), and Mitochondrial Rho GTPases (MIROs)/RHOTs leading to its further recruitment and retention at the MOM via a Ser65-phosphorylated-ubiquitin (phospho-ubiquitin) dependent feed-forward amplification mechanism (Ordureau et al., 2014, Ordureau et al., 2020, Ordureau et al., 2018, Sarraf et al., 2013, Chan et al., 2011, Narendra et al., 2012). The presence of ubiquitin and/or phospho-ubiquitin on damaged mitochondria stimulates recruitment of autophagy adaptor receptors of which NDP52, OPTN and to a small extent Tax1bp1 are required for efficient clearance of damaged mitochondria (Lazarou et al., 2015, Ordureau et al., 2018, Heo et al., 2015).

The initial mechanisms by which Parkin is activated and identification of bona fide substrates has been mainly studied in human cancer lines over-expressing Parkin and to date very few studies have assessed its molecular function in physiologically relevant neuron cell types at the endogenous level. Previous analysis of ubiquitylation in HeLa cells over-expressing Parkin found highest levels of ubiquitylation in the abundant VDAC1/2/3 proteins and also revealed moderate ubiquitylation of specific sites within MFN2, CISD1 and TOMM20 proteins (Sarraf et al., 2013, Rose et al., 2016, Ordureau et al., 2018). Ubiquitylation analysis in human ESC derived iNeurons (expressing markers of excitatory cortical neurons) has revealed endogenous-PINK1-dependent accumulation of phospho-ubiquitin; a 70, 14 and 2 fold increase in K63, K11 and K6 ubiquitin linkages (the latter being lower than that observed in HeLa cells over-expressing Parkin); and 134 ubiquitylation sites spanning 83 proteins of which the majority are localised at the MOM including VDAC1/3 and MFN2 (Ordureau et al., 2018, Ordureau et al., 2020). Collectively these studies have elaborated a model which suggests that PINK1 and Parkin dependent accumulation of ubiquitin chain types mainly at the MOM is sufficient for downstream signalling and mitochondrial clearance (Ordureau et al., 2020, Swatek et al., 2019).

In previous studies we have reported that endogenous PINK1 and Parkin activation can be robustly measured in primary cortical neurons derived from mouse embryonic-derived neuronal progenitors, that is an established and widely validated cell system to investigate neurobiological mechanisms relevant to mature neurons (Gordon et al., 2013, Barini et al., 2018, McWilliams et al., 2018). Herein, we have undertaken global diGLY ubiquitylation analysis in primary cortical neurons of wild-type and homozygous PINK1 knockout mice to map the ubiquitin architecture of mitochondria under normal basal conditions and upon activation of PINK1 upon mitochondrial depolarisation. After 5 h of stimulation we detect ubiquitylation of 58 mitochondrial proteins located mainly within the MOM; associated with an increase in K63 ubiquitin chain linkage type; and a low level of mitochondrial protein turnover at this timepoint that is blocked in PINK1 knockout neurons. We further demonstrate the regulation of mitochondrial substrates by Parkin using cell-based assays in Parkin knockout neurons and Parkin reconstitution *in vitro* catalytic activity assays. Comparative analysis with human iNeurons datasets elaborated a PINK1/Parkin-dependent ubiquitylation signature on depolarised mitochondria in neurons comprising 49 sites across 22 MOM proteins conserved between mouse and human neurons that represents a cellular readout for PINK1 and Parkin activity as well as a potential proxy readout of Parkinson’s disease linked mitochondrial dysfunction in future studies.

## RESULTS

### Characterisation of PINK1-induced ubiquitin signalling in mouse neurons

We initially characterised PINK1 and Parkin expression and activity in primary cortical neuron cultures established from E16.5 mouse embryos at 21 days *in vitro* (DIV) (Figure 1A). We undertook a timecourse analysis of PINK1 activation following Antimycin A (10μM) / Oligomycin (1μM) stimulation (inhibition of respiratory chain enzyme complex II/III and ATP synthase respectively; AO). Endogenous PINK1 levels were detectable following immunoprecipitation-immunoblot analysis revealing low expression in neurons under basal conditions and increased expression over time upon mitochondrial depolarisation (Figure S1A). Immunoblot analysis of Parkin demonstrated basal expression that was stable upon mitochondrial depolarisation up to 9h (Figure S1A). We optimised the detection of phospho-ubiquitin using HALO-multiDSK and immunoblotting with anti-phospho-Ser^65^ Ubiquitin antibody and signal was abolished using a mutant form of HALO-multiDSK (Figure S1B). Furthermore, we did not observe any difference in the detection of phospho-ubiquitin using Tandem Ubiquitin Binding Entities (TUBE) pulldowns with HALO-multiDSK and HALO-UBA^UBQLN1^ (Figure S1B). Upon mitochondrial depolarisation we observed robust accumulation of phospho-ubiquitin at 3 h and this was maximal at 5 h and similarly we also detected phospho-Parkin upon mitochondrial depolarisation (Figure S1A). We therefore undertook all subsequent analysis at this timepoint. We next investigated signalling in PINK1 knockout neurons and observed complete loss of phospho-ubiquitin (Figure 1B). We have previously found that PINK1 activation leads to induction of phosphorylation of a subset of Rab GTPases including Rab8A at Serine 111 (Ser111) in human cancer cell lines (Lai et al., 2015) and upon mitochondrial depolarisation we observed Ser111-phosphorylated Rab8A (phospho-Rab8A) in wild-type neurons and this was abolished in PINK1 KO neurons (Figure 1B). Upon mitochondrial depolarisation of wild-type and Parkin KO neurons, we observed loss of phospho-Parkin in Parkin KO neurons and, whilst the level of phospho-ubiquitin was substantially reduced, it was not abolished suggesting that other unknown ubiquitin E3 ligases may contribute ubiquitin marks that can be phosphorylated by PINK1 in mature neurons (Figure S1C), a finding consistent with studies in Parkin-deficient iNeurons (Ordureau et al., 2018). Furthermore we observed decreased PINK1 levels in Parkin KO neurons consistent with previous observations in human Parkin S65A iNeurons (Ordureau et al., 2018) and human Parkin S65N patient derived fibroblasts (McWilliams et al., 2018) suggesting a potential feedback mechanism of Parkin activation on PINK1 stabilisation. We have previously observed that PINK1-induced phospho-Rab8A can occur in HeLa cells that lack Parkin or in Parkin KO mouse embryonic fibroblasts (MEFs) and consistent with this, we did not observe any difference in phospho-Rab8A in wild-type or Parkin KO neurons following mitochondrial depolarisation (Figure S1C). It has been reported that additional monogenic forms of Parkinson’s may interplay with the PINK1/Parkin pathway including the Asp620Asn (D620N) mutation of the retromer associated VPS35 gene (Williams et al., 2018). However, we did not observe any alteration in phospho-ubiquitin or phospho-Rab8A in VPS35 D620N homozygous and heterozygous neurons compared to wild-type controls following mitochondrial depolarisation (Figure S2).

**Figure 1.**
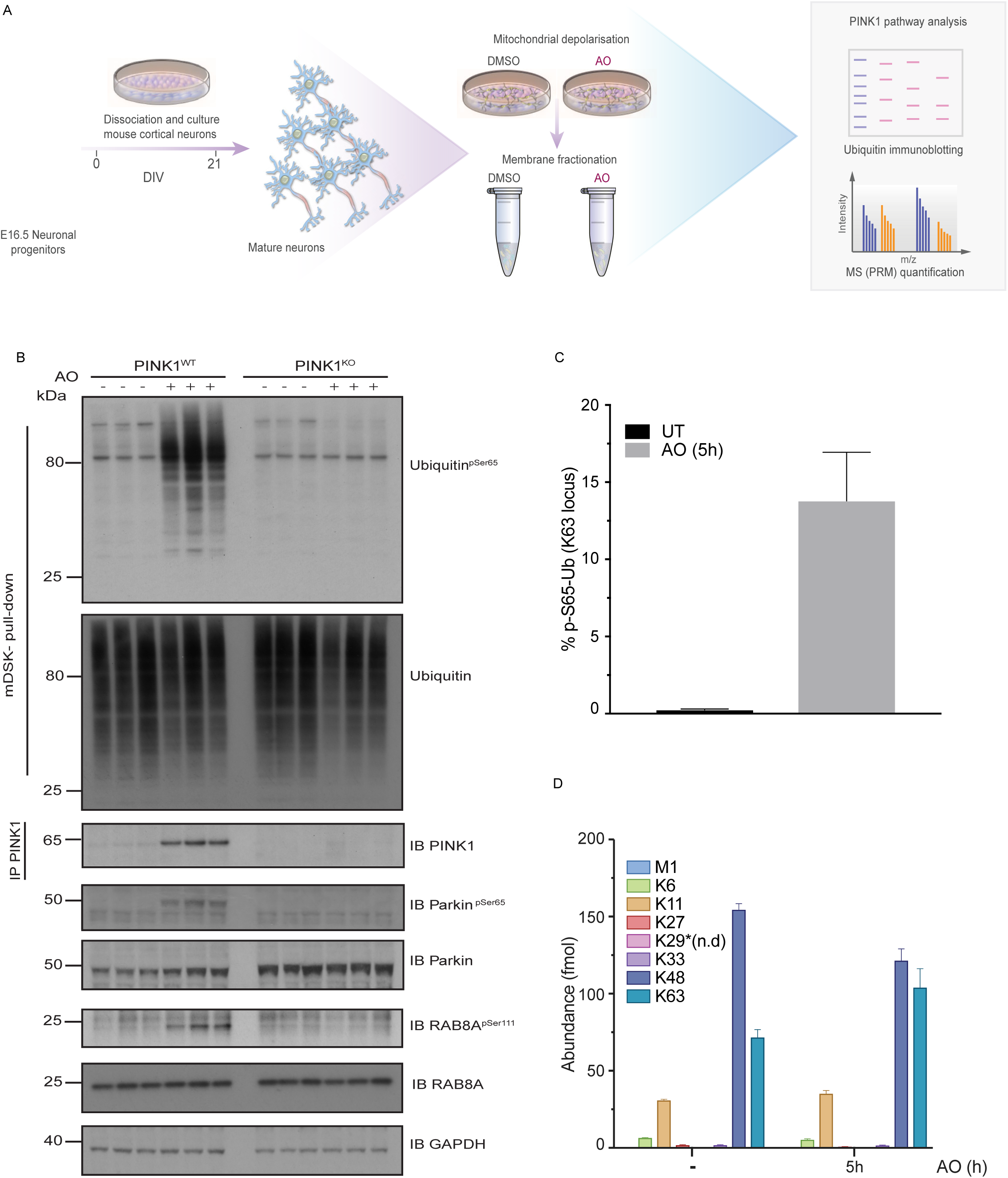
PINK1 signalling in mouse cortical neurons. **A.** Experimental workflow in primary mouse neurons. E 16.5 cortical neurons were cultured for 21 days in vitro (DIV) and membrane enrichment was performed, after mitochondrial depolarisation induced with 10 μM of Antimycin A combined with 1 μM of Oligomycin for 5 h. **B.** Immunoblots showing comparative analysis of phospho-Ser65 Ubiquitin levels in primary cortical neuron cultures from wild-type and PINK1 knockout (KO) mice. Cultures were stimulated with Antimycin A and Oligomycin for 5 h prior to membrane enrichment. Phospho-Ser65 Ubiquitin was detected by immunoblotting after ubiquitin enrichment by incubating with ubiquitin-binding resin derived from Halo-multiDSK (mDSK). Affinity captured lysates were also subjected to immunoblotting with total ubiquitin antibody. Immunoprecipitation (IP)-immunoblot showed PINK1 protein stabilisation after mitochondrial depolarisation. Phospho-Ser111 Rab8A and phospho-Ser65 Parkin were detected by immunoblotting. Lysates were also subjected to immunoblotting with indicated antibodies for loading and protein expression controls. **C and D**. C56BL/6J mouse cortical neurons (DIV 21) were depolarized with AO for 5 hours and whole cell lysates were subjected to Pt-PRM quantification. Abundance (fmol) for individual Ub chain linkage types. UT untreated (D) and percentage of phospho-Ser65 Ubiquitin (C) is plotted. Error bars represent SEM (n = 3). n.d., not determined.

We next employed targeted Parallel Reaction Monitoring (PRM) proteomic analysis in whole cell lysates of mature DIV21 neurons to quantitatively assess ubiquitin changes upon AO stimulation and observed an approximate 52 fold increase in phospho-ubiquitin with a stoichiometry of ubiquitin phosphorylation of ∼0.13 at 5 h treatment (Figure 1C). Under basal conditions we detected nearly all ubiquitin chain linkage types including K6, K11, K27, K33, K48 and K63 and upon 5 h of AO treatment, we only observed an increase in K63 chain linkages (Figure 1D). In parallel studies, we undertook PRM analysis of less mature DIV12 neurons in which Parkin expression is lower (Barini et al., 2018) and stimulated these cells with AO for less time at 3 h. Interestingly in TUBE pulldowns of whole cell lysates or mitochondrial-enriched fractions, we observed an approximate 300-fold and 40-fold increase in phospho-ubiquitin respectively that was not associated with any significant increase in ubiquitin chain linkage types (Figure S3). This suggests that basal mitochondrial ubiquitin levels in neurons may be sufficient to promote the initial generation of phospho-ubiquitin under conditions where Parkin activity is not maximal.

### Quantitative proteome and diGLY proteome of mitochondria in mouse cortical neurons under basal conditions and mitochondrial depolarisation

We next utilised a tandem mass tagging (TMT)-MS3-based pipeline (Rose et al., 2016) (Figure 2A) to quantify the mitochondrial proteome abundance and the mitochondrial ubiquitylome under basal conditions and upon mitochondrial depolarization that has been previously deployed in human iNeurons (Ordureau et al., 2020). We isolated mitochondria-containing membrane fractions from quintuplicate cultures of DIV21 C57BL/6J mouse primary cortical neurons that were untreated or treated with 10μM AntimycinA and 1μM Oligomycin (AO) for 5 h (Figure S4A-B). We observed few changes in the proteome abundance following mitochondrial depolarisation; of the 6255 proteins quantified (Table S1) 4 proteins (CLSTN2, SYT5, MKL1, HIST3H2BA) were significantly reduced and one protein (PPM1H) significantly increased after AO treatment (Figure S4C). There was a slight shift to the left in the AO proteome compared to the untreated proteome suggesting a low level of turnover (Figure S4C). This is consistent with analysis undertaken in iNeurons stimulated with AO (Ordureau et al., 2020).

**Figure 2.**
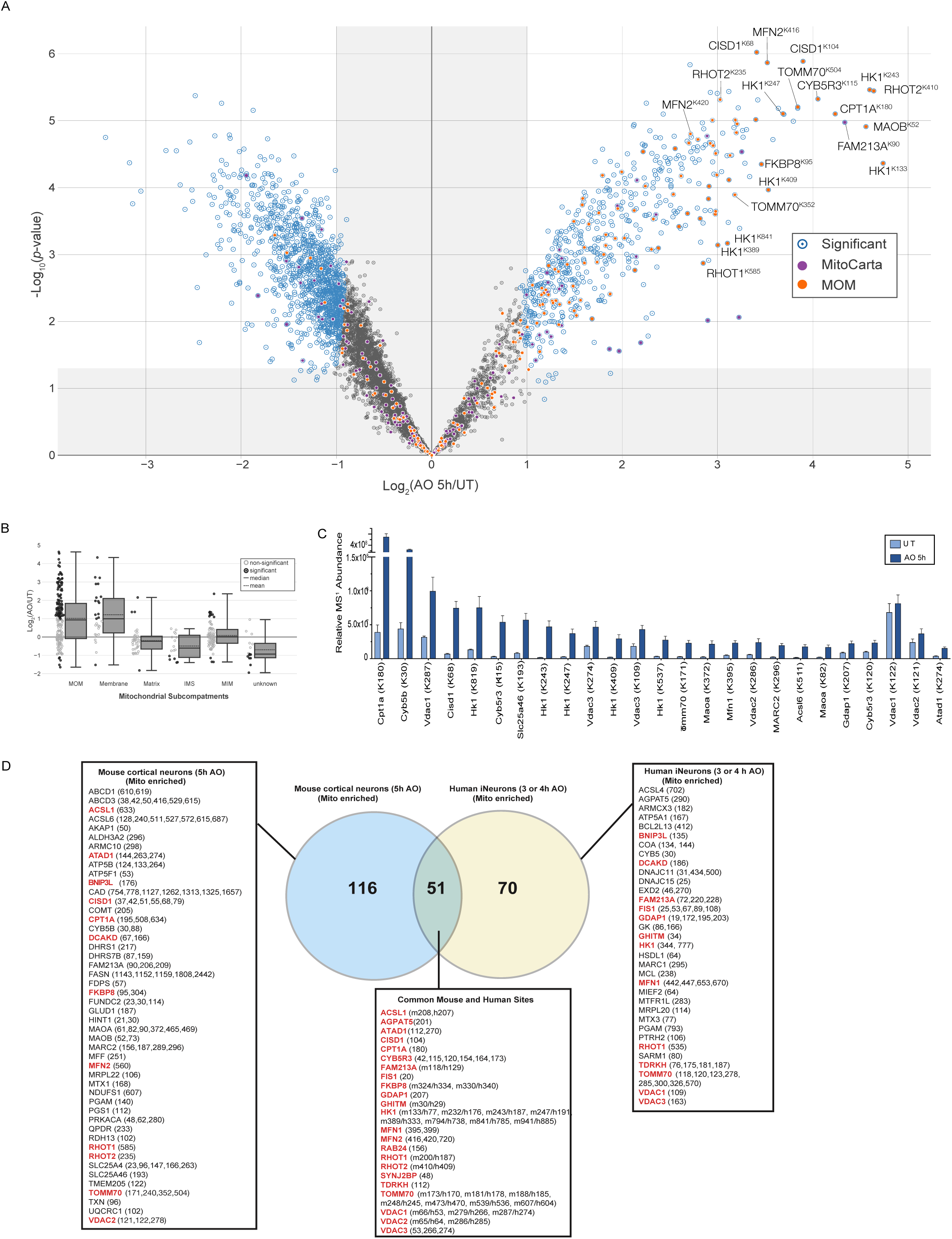
Global ubiquitylation analysis of mitochondria in neurons upon mitochondrial depolarisation. **A.** C56BL/6J primary cortical neurons were depolarized with AO (5 h) and membrane enriched lysates were subjected to diGLY capture proteomics. Fold increase for individual ubiquitylated targets is shown in the Volcano plot. The x-axis specifies the fold-changes (FC) and the y-axis specifies the negative logarithm to the base 10 of the t-test p-values. 1606 dots reflect the significant hits (Welch’s t-test (S0=2), corrected for multiple comparison by permutation-based FDR (1%))). 559 and 1047 dots respectively represent ubiquitylated targets upregulated or downregulated after mitochondrial depolarisation. diGLY-peptide of proteins associated with mitochondria (MitoCarta 3.0) or Mitochondrial outer membrane localization are indicated. **B.** Distribution of changes in diGLY peptides for proteins that localize in the mitochondrial subcompartments: matrix, MIM, MOM, IMS, membrane and unknown (Mitocarta 3.0). **C.** Ranking analysis of top mito Ub sites – MS1- and TMT-based intensity of all diGLY peptides was extracted and the top 25 diGLY sites with the largest relative abundance change upon 5h depolarization is indicated. Error bars represent SEM (n = 5). **D.** Venn diagram of overlapping diGLY sites observed in site observed for mitochondrial enriched mouse cortical neurons (5 h post-depolarization) and sites observed from mitochondrial enriched human iNeurons (3 or 4 h post depolarisation) (Ordureau et al., 2020). All peptides used were increased by at least 2-fold (log_2_ ratio > 1.0), with p < 0.05.

We next determined the mitochondrial ubiquitylome in mouse neurons using diGLY affinity capture coupled with quantitative proteomics (Figure 2A, Figure S4-B). Tryptic peptides from membrane enriched extracts of mouse primary cortical neurons untreated or treated with AO for 5 h were subjected to α-diGLY immunoprecipitation and samples analysed using 11-plex TMT-MS^3^ with diGLY peptide intensities normalised with total protein abundance measured in parallel (Figures 2A; Table S1; see STAR Methods). We detected and quantified ∼9154 diGLY-containing Kgg peptides of which 5616 are unique sites. Of those unique sites, 559 are statistically up-regulated (normalised for total proteome) upon AO treatment (Figure 2A). We next analysed the data for the number of ubiquitylated mitochondrial proteins using the recently published Mitocarta 3.0 database containing data on 1140 mouse gene encoding proteins that are strongly localised to mitochondria including their sub-mitochondrial location (Rath et al., 2021). 345 unique diGLY-containing Kgg peptides were detected in a total of 139 mitochondrial proteins under basal and induced conditions of which 54 proteins were located in the mitochondrial outer membrane (MOM); 42 proteins in the mitochondrial inner membrane (MIM), 19 proteins in the matrix; 10 proteins of unknown mitochondrial sublocation; 9 proteins that are membrane associated and 5 proteins in the intermembranous space (IMS) (Figure 2B). The major portion of ubiquitylated proteins up-regulated by AO were MOM localised accounting for 16.6% (MitoCarta 21.3%) after 5 h of mitochondrial depolarization (Figure 2B). We also detected Kgg peptides of several mitochondrial membrane proteins, TDRKH, ABCD3, DCAKD, DHRS7B; MIM proteins RDH13, PGS1 and matrix proteins, GLUD1, MRPL22 that were increased ∼2 fold or more following AO (Table S1). Based on MS1-abundance information and TMT quantification, we performed a simple ranking analysis of Kgg peptides cross-referenced to Mitocarta 3.0 (Figure 2C) to determine the most abundant MOM ubiquitylation sites and these included CPT1α (K180, K195, K508); CYB5B (K30); VDAC1 (K122, K187); HK1 (K819, K243, K247), CISD1 (K68, K79), CYB5R3 (K115, K120) and TOMM70 (K171, K240) (Figure 2C). Conversely we also performed similar analysis on MOM ubiquitylation sites that are most reduced following AO treatment (Figure S5A). This revealed down-regulated sites for PARK7/DJ1 (K93), VDAC1 (K33), PGAM5 (K73, K140, K161), and TOMM20 (K56, K61) (Figure S5A). Comparison of the basal ubiquitin levels of the most up-regulated and down-regulated MOM sites did not generally predict their subsequent modification following AO treatment; for example HK1, CYB5R3 and TOMM70 exhibited basal ubiquitin at sites in the range of 1.5 x10^6^ but became highly modified (Figure 2C and Figure S5A).

Overall 167 Kgg peptides from 60 mitochondrial proteins were significantly elevated by AO treatment (Figure 2D and Figure S5B) and this compared to 121 Kgg peptides from 46 mitochondrial proteins that were significantly elevated by AO treatment in human iNeurons (Ordureau et al., 2020).

Consistent with previous analysis in human iNeurons, we found that the majority of proteins were MOM localised and 51 ubiquitylation sites spanning 23 mitochondrial proteins were conserved between mouse neurons and human iNeurons namely ACSL1, AGPAT5, ATAD1, CISD1, CPT1α, CYB5R3, FAM213A, FIS1, FKBP8, GDAP1, GHITM, HK1, MFN1/2, RAB24, RHOT1/2, SYNJ2BP, TDRKH, TOMM70, and VDAC1/2/3 (Figure 2D and Figure S5B). In addition we identified two proteins whose ubiquitylation was increased in both mouse neurons and human iNeurons although the sites were distinct namely, the mitophagy receptor BNIP3L, and DCAKD (Figure 2D). Of mitochondrial substrates specific to mouse neurons, we interestingly detected ubiquitylation of enzymes linked to dopamine and catecholamine metabolism including Monoamine Oxidase A (MAO-A), Monoamine Oxidase B (MAO-B) and Catechol O Methyltransferase (COMT) which degrades dopamine and are established drug targets for symptomatic treatment of Parkinson’s (Table S1).

Previous TMT analysis of HeLa cells (over-expressing Parkin) has indicated that activation of PINK1 and Parkin may stimulate recruitment of a large panoply of proteins to depolarized/damaged mitochondria as determined by the detection of 137 ubiquitylation sites derived from 85 cytosolic proteins including Parkin and autophagy receptors such as OPTN and TAX1BP1 in mitochondrial-enriched fractions (Rose et al., 2016). Under endogenous conditions we were unable to detect ubiquitylation sites for Parkin or autophagy receptors in our analysis but could detect unmodified peptides for OPTN and TAXBP1 (Table S1). Comparative analysis of datasets revealed approximately 80 cytosolic proteins whose ubiquitylation was increased upon AO treatment in both mouse neuron and HeLa cell analyses and in a significant number of proteins, the site of ubiquitylation was identical between datasets (Table S2). These common sites included the p97 ubiquitin-binding co-factor FAF2 (K167); the ALS-linked protein TDP-43 (K181), the protein kinase AKT1 (K426) and SUMO E3 ligase, TRIM28 (K320) (Table S2) (Rose et al., 2016). Interestingly in neurons we also found a large number of ubiquitylation sites in cytosolic kinases that were up-regulated upon AO treatment including GSK3β, CAMK2α/β/γ several PRKC isoforms, ULK1 (K162) and LRRK2 (K1132). Recent studies have indicated a role for ULK1 in the initiation of mitophagy (Vargas et al., 2019) and LRRK2 has also been shown to regulate basal mitophagy (Singh et al., 2021) and in future work it will be interesting to determine whether other cytosolic kinase and phosphatase signalling components identified in our screen may be implicated in mitophagy mechanisms (Table S3).

### Quantitative diGLY proteome analysis in mitochondria of wild-type and PINK1 knockout mouse cortical neurons

We next determined which ubiquitylated substrates were dependent on activation of PINK1 (Figure S6A). We isolated mitochondria from cultures of primary cortical neurons from crosses of wild-type and PINK1 knockout mice that were untreated or treated with AO for 5 h (Figures S6A-B). We compared the proteome of triplicate cultures of WT and PINK1 KO mouse cortical neurons that were untreated or treated with AO for 5 h. Similar to the previous experiment we found very little change in the proteome abundance of wild-type neurons and the total proteome was slightly reduced in wild-type neurons following AO treatment indicating increased turnover (Figure 3A and S6C). Interestingly this mild reduction was completely blocked in the PINK1 knockout neurons (Figure3A and S6C). These results are consistent with similar observation of the proteome of wild-type and Parkin S65A iNeurons following AO treatment for 6 h (Ordureau et al., 2020). Furthermore, quantitative TMT analysis demonstrated a ∼50-fold increase in Ser65-phospho-ubiquitin upon AO in wild-type neurons that was abolished in PINK1 knockout neurons (Figure 3B). In parallel we also measured Ser57-phospho-ubiquitin that is independent of AO treatment and there was no difference between wild-type and PINK1 KO neurons (Figure 3B).

**Figure 3.**
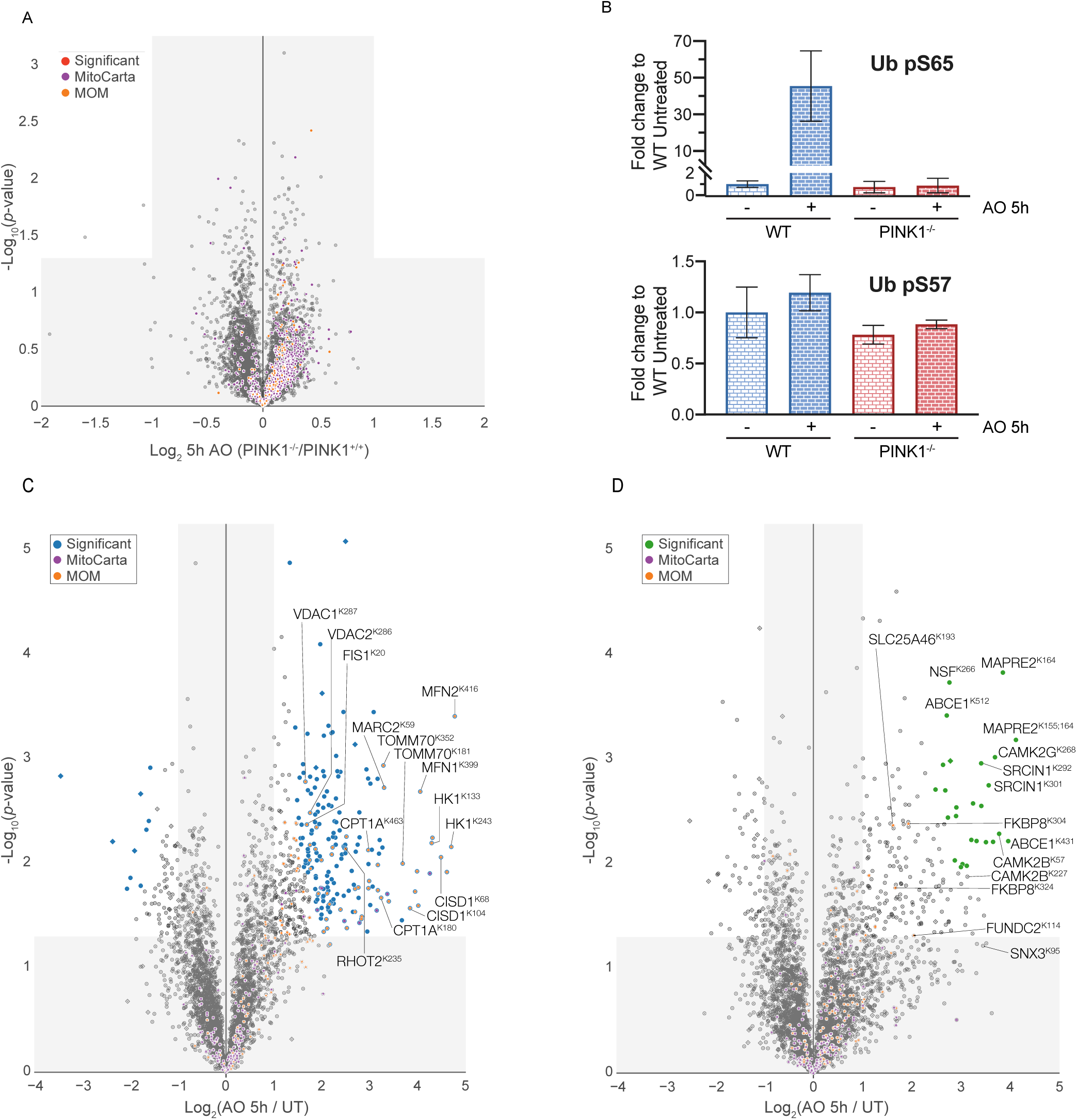
Global ubiquitylation analysis of mitochondria in neurons of PINK1 WT and KO neurons. **A.** Total protein abundance in wild-type or PINK1 deficient neurons. Membrane enriched lysates from C56BL/6J primary cortical neurons were subjected to quantitative proteomics. Fold increase for individual protein is shown in the Volcano plot. The x-axis specifies the fold-changes (FC) and the y-axis specifies the negative logarithm to the base 10 of the t-test p-values (Welch’s t-test (S0=0.585), corrected for multiple comparison by permutation-based FDR (5%)). Proteins associated with mitochondria (MitoCarta 3.0) or Mitochondrial outer membrane localization are indicated. **B.** Abundance for phosphorylated-Serine 65 (upper panel) or-Serine 57 (lower panel) of Ub was quantified and plotted as fold change to untreated sild-type samples. C) is plotted. Error bars represent SEM (n = 3). n.d., not determined. Error bars represent SEM (n = 3, 3, 3, 2). **C.** Same as (A) but membrane enriched lysates derived from wild-type cells were subjected to diGly capture proteomics. Fold increase for individual ubiquitylated targets is shown in the Volcano plot. The x-axis specifies the fold-changes (FC) and the y-axis specifies the negative logarithm to the base 10 of the t-test p-values. 198 dots reflect the significant hits (Welch’s t-test (S0=1), corrected for multiple comparison by permutation-based FDR (1%)). 187 and 11 sites respectively represent ubiquitylated targets upregulated or downregulated after mitochondrial depolarisation. diGLY-peptide of proteins associated with mitochondria (MitoCarta 3.0) or Mitochondrial outer membrane localization are indicated. Square shaped dot, indicate a ubiquitylated peptide that was not normalized to its protein abundance (not determined). **D.** Same as (A) but membrane enriched lysates derived from PINK1-deficient cells were subjected to diGly capture proteomics. 26 dots reflect the significant hits (Welch’s t-test (S0=1), corrected for multiple comparison by permutation-based FDR (1%)). 26 and 0 sites respectively represent ubiquitylated targets upregulated or downregulated after mitochondrial depolarisation. diGLY-peptide of proteins associated with mitochondria (MitoCarta 3.0) or Mitochondrial outer membrane localization are indicated. Square shaped dot, indicate a ubiquitylated peptide that was not normalized to its protein abundance (not determined).

We next determined the mitochondrial ubiquitylome in mouse neurons using diGLY affinity capture coupled with quantitative proteomics (Figures S6A-B) (Ordureau et al 2020). Tryptic peptides from mitochondrial enriched extracts of primary cortical neurons of wild-type and PINK1 knockout mice with or without mitochondrial depolarisation (5 h) were subjected to α-diGLY immunoprecipitation and samples analysed using 11-plex TMT-MS^3^ (WT: 3 untreated/3AO; KO: 3 untreated/2AO) with diGLY peptide intensities normalised with total protein abundance measured in parallel (Figures 3C-D and Figures S6A – B) (Table S4) (see STAR Methods).

We quantified a total of ∼7210 diGLY-containing Kgg peptides of which ∼3951 are unique sites (Figure 3C). From these, we identified 177 ubiquitylation sites in 99 proteins in wild-type neurons after 5 h of AO stimulation whose abundance was significantly increased (Figure 3C) and the majority of sites had been detected in AO treated C57BL/6J neurons (Figure 2). AO treatment of PINK1 KO mouse neurons for 5 h led to substantially reduced AO-induced ubiquitylation of mitochondrial proteins including all but one of the common set of 23 MOM proteins (FKBP8) identified previously (Figure 3D and S6D). In contrast we observed ubiquitylation of a number of cytosolic proteins including CAMK2α/β/γ, and SNX3 that whose ubiquitylation was independent of PINK1 (Figure 3D and S6D). Principal-component analysis revealed that the 5 h depolarization data of WT was not similar to the untreated WT and or PINK1 KO samples, which were also more similar to each other (Figure S6E).

### Validation of Parkin-dependent substrates using cell-based and *in vitro* studies

Based on the ranking of the most abundant ubiquitylated (Kgg) mitochondrial sites (Figure 2A, 2C and Figure S5B), we proceeded to investigate whether we could detect endogenous substrate ubiquitylation via biochemical analysis in wild-type C57BL/6J neurons that were either untreated or stimulated with AO for 5 h. Mitochondrial-enriched fractions from neurons were subjected to TUBE pulldowns with HALO-UBA^UBQLN1^ or HALO-multiDSK (and non-binding multiDSK mutant) and immunoblotted with specific antibodies for each substrate as well as phospho-ubiquitin to confirm Parkin activation (Figure S7). This confirmed previous observations (Barini et al., 2018, McWilliams et al., 2018) that CISD1 can undergo multi-monoubiquitylation upon AO treatment and the signal was similar for HALO-UBA^UBQLN1^ or HALO-multiDSK pulldowns. We also observed robust monoubiquitylation of CPT1α, an integral outer membrane protein that converts activated fatty acids into acylcarnitines and facilitates their transport into mitochondria (Lee et al., 2011) (Figure S7). This was more easily detected with HALO-multiDSK pulldown and we employed HALO-multiDSK to undertake a timecourse analysis and confirmed that CPT1α undergoes time-dependent ubiquitylation similar to CISD1 following AO treatment (Figure 4A). Interestingly we observed a high molecular weight species of ∼140 kDa that was immunoreactive to the CPT1α antibody following HALO-mDSK pulldown (Figure 4A, S7-S8). This was present under basal conditions but increased after AO treatment. CPT1α is reported to form complexes at the MOM as well as oligomerise (Faye et al., 2007) and we cannot rule out that the upper band represents oligomerised species of ubiquitylated CPT1α. The detection of AO induced ubiquitylation of other mitochondrial proteins varied depending on the ubiquitin-affinity method used including TDRKH and ATAD1 that were selectively detected via HALO-UBA^UBQLN1^ pulldown (Figure S7). There were also several mitochondrial proteins exhibiting substantial basal ubiquitylation including AGPAT5, MFN2, and TOMM70, that prevented unambiguous detection of enhanced ubiquitylation upon AO treatment by our TUBE-based assay (Figure S7).

**Figure 4.**
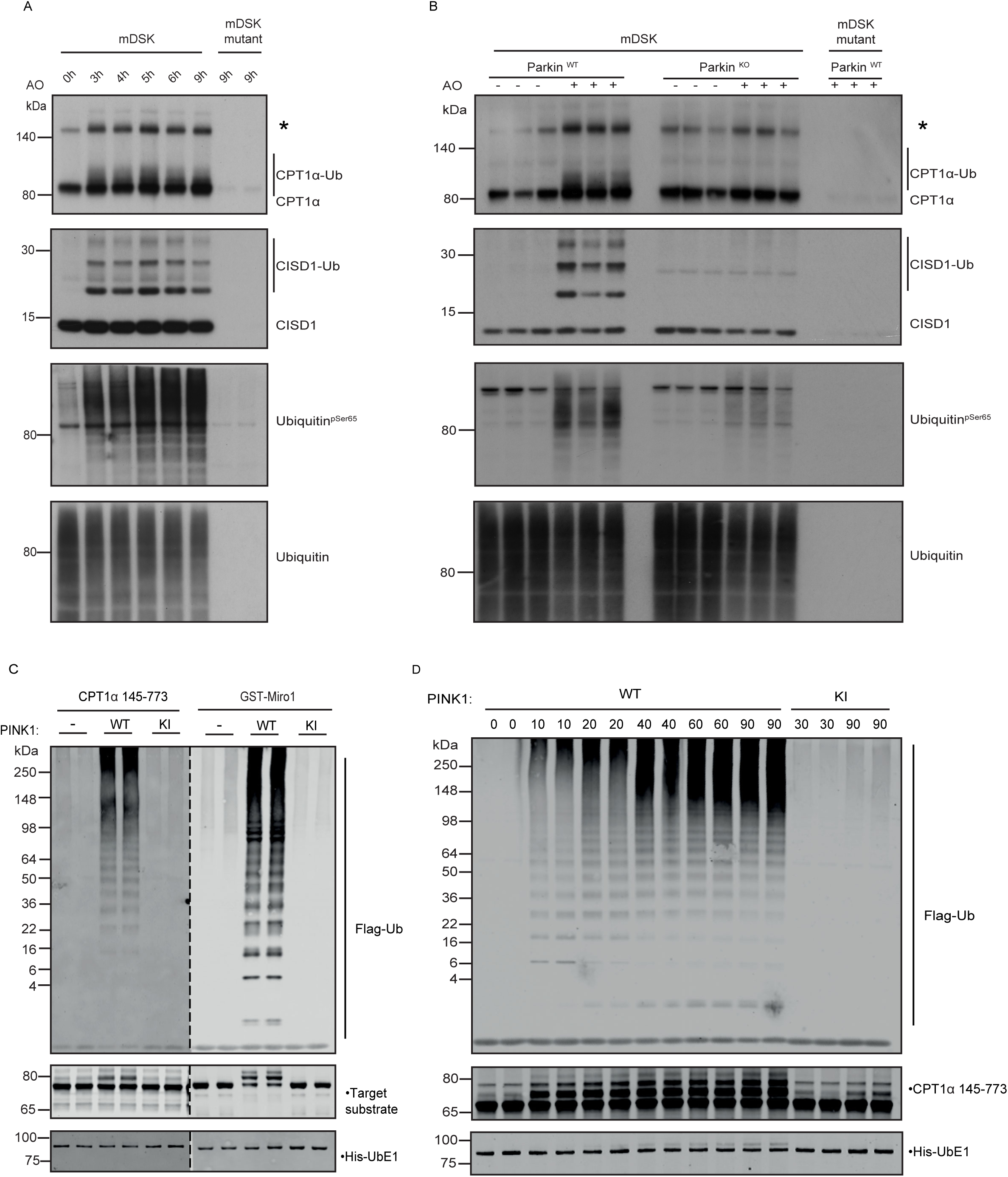
Validation of CPT1α as a Parkin substrate. **A.** Time-course analysis of CPT1α and CISD1 ubiquitylation following AO stimulation in C57BL/6J cortical neurons. Ubiquitylation detection by Halo-multiDSK (mDSK) pull-down prior to immunoblotting with anti-CPT1α, anti-CISD1, anti-phosphoSer65 Ubiquitin and anti-Ubiquitin antibodies. * High molecular weight CPT1α reactive species. **B.** CPT1α ubiquitylation is abrogated in Parkin KO neurons. Membrane lysates of Parkin WT and KO cortical neurons after 5 hours of AO stimulation, were subjected to ubiquitylated-protein capture by Halo-multiDSK (mDSK), prior to immunoblot with anti-CPT1α, anti-CISD1, anti-phosphoSer65 Ubiquitin and anti-Ubiquitin antibodies. * High molecular weight CPT1α reactive species. **C and D.** Parkin ubiquitylates CPT1α *in vitro.* C. Recombinant CPT1α protein was assessed for ubiquitylation by Parkin. Miro1 recombinant protein was used as positive control. D. Timecourse for CPT1α ubiquitylation by Parkin. For ubiquitylation assays Parkin was first activated using a 30 min kinase reaction at 37 °C in the presence or absence of 0.36 μM wild type (WT) or kinase inactive (KI) TcPink1. Ubiquitylation reactions were started by the addition of 0.5 μM the target protein, 50 μM Flag-Ubiquitin, 0.12 μM His-UbE1 and 1 μM UbE2L3. Ubiquitylation reactions were stopped after a 30 min incubation at 37 °C unless stated otherwise by the addition of 4 x LDS loading buffer. 0.625 pmol target protein was resolved on a 4 to 12 % Bis-Tris gels, transferred to nitrocellulose membranes, and immunoblotted using anti-Flag, anti-His, and anti-targeted protein respectively. Membranes were imaged using LiCor.

We next assessed CPT1α ubiquitylation alongside CISD1 in triplicate biological replicates of Parkin knockout (Figure 4B) and PINK1 knockout neurons (Figure S8A) with their respective wild-type littermate controls. Mitochondrial fractions from neurons untreated or stimulated with AO for 5 h were subjected to TUBE pulldowns with HALO-multiDSK demonstrating that CPT1α monoubiquitylation was abolished in Parkin and PINK1 knockout neurons and this paralleled loss of CISD1 ubiquitylation (Figure 4B and Figure S8). Furthermore, the increased ubiquitylation of the ∼140kDa CPT1α-immunoreactive species was also attenuated in PINK1 and Parkin knockout neurons. We also explored whether CPT1α ubiquitylation can independently be detected in a different cell system and analysed human SH-SY5Y neuroblastoma cell lines that express endogenous PINK1 and Parkin (Figure S8B). Upon addition of AO, we observed time-dependent accumulation of CPT1α ubiquitylation that was maximal at 9 h associated with an increase in endogenous phospho-Parkin (Figure S8B). From 9 – 24 h, the levels of total Parkin and phospho-Parkin significantly reduced and this was accompanied by a decrease in CPT1α ubiquitylation consistent with its Parkin-dependence (Figure S8B).

We have previously elaborated a robust *in vitro* assay of Parkin E3 ligase activity that monitors multi-monoubiquitylation of RHOT1/Miro1 (residues 1-592) as well as the formation of free poly-ubiquitin chains (Kazlauskaite et al., 2014a). To assess whether Parkin (activated by phosphorylation of PINK1) directly ubiquitylates CPT1α, we phosphorylated untagged full-length Parkin with the active insect orthologue of PINK1, Tribolium castaneum (TcPINK1), in the presence of adenosine triphosphate (ATP) and then added a reaction mix containing E1 ubiquitin-activating ligase, UbcH7 conjugating E2 ligase, ubiquitin, Mg-ATP and a fragment of CPT1α (residues 145-773) alongside GST-Miro1 (residues 1-592) that were expressed and purified in *E. coli* (Figure S9). After 30 min, reactions were terminated with SDS sample buffer in the presence of 2-mercaptoethanol and heated at 100 ^°^C, substrate ubiquitylation was assessed by immunoblot analysis with antibodies that detect ubiquitin (FLAG), CPT1α, and Miro1. Consistent with previous findings in the absence of PINK1 phosphorylation, Parkin was inactive as no evidence of free ubiquitin chain formation or Miro1 ubiquitylation was observed; with the addition of wild-type TcPINK1, Miro1 multi-monoubiquitylation in addition to free polyubiquitin chain formation was observed (Figure 4C). No significant Miro1 ubiquitylation or polyubiquitin chain formation was observed in the presence of the kinase-inactive TcPINK1 (D359A) (Figure 4C). Under these conditions we observed multi-monoubiquitylation of CPT1α indicating that this is also a direct substrate of Parkin. To determine the temporal dynamics of CPT1α ubiquitylation, we undertook a timecourse analysis and observed the appearance of multi-monoubiquitylation at 10 min following addition of activated Parkin and this increased in a time-dependent manner and paralleled polyubiquitin chain formation (Figure 4D). We also tested additional mitochondrial substrates identified from our diGLY mass spectrometry screen including FAM213A, MAO-A, MAO-B and Acyl-CoA Synthetase Long Chain 1 (ACSL) (ACSL1; residues 46-end) that forms a complex with CPT1α (and VDAC) to promote influx of activated fatty acids into the mitochondria (Lee et al., 2011) (Figure 2D and S9). All underwent mono-ubiquitylation following addition of PINK1-activated Parkin further suggesting that Parkin preferentially catalyses mono/multi-monoubiquitylation of substrates *in vitro* (Figure S9 and S10).

We next investigated whether non-mitochondrial proteins whose ubiquitylation was increased following AO treatment are direct Parkin substrates (Figure 2A, 3D, and S11A). Under similar conditions described above, we incubated SNX3, CAMK2 α and CAMK2β with activated Parkin *in vitro* (Figure S11B-D, Left panel). We observed free polyubiquitin chain formation but no significant substrate phosphorylation indicating that these are not Parkin substrates and unlikely to be regulated by active Parkin in neurons. Mass spectrometry data did not reveal any difference in sites of ubiquitylation in these proteins between wild-type and PINK1 knockout neurons following stimulation with AO suggesting that additional ubiquitin E3 ligases may be activated and recruited to the mitochondria upon AO treatment in a PINK1 independent manner (Figure 3D, Figure S11B-D, Right panel).

## DISCUSSION

Our groups have previously demonstrated the utility of primary mouse neurons and human iNeurons to study endogenous Parkin activation including conservation of Parkin activation by PINK1-dependent phosphorylation of Serine65 (McWilliams et al., 2018, Ordureau et al., 2020, Ordureau et al., 2018). Combining a diGLY/TMT-MS^3^ workflow (Rose et al., 2016) in mouse neurons with comparative data from previous analysis in human iNeurons (Ordureau et al., 2020) has elaborated a defined set of physiological substrates of Parkin conserved in mouse and human neuron cell types (Figure 2). Strikingly from many thousands of peptides detected by mass spectrometry in the screen, we found 49 unique diGLY sites in 22 proteins in which ubiquitylation was increased greater than 2-fold or more in neurons upon activation of PINK1 by mitochondrial depolarisation (Figure 2 and 3). We also identified 2 proteins, BNIP3L and DCAKD, whose ubiquitylation was increased in both mouse and human iNeurons albeit at different sites (Figure 2). Furthermore, all of the sites were present on proteins localised at the mitochondrial outer membrane (MOM) consistent with previous studies on how activation of PINK1 and Parkin on the MOM is sufficient to induce clearance of damaged mitochondria via mitophagy (Harper et al., 2018, Singh and Muqit, 2020, Moehlman and Youle, 2020). This was associated with the enhanced formation of phospho-ubiquitin and K63 chain linkages in mouse neuronal mitochondria (Figure 1D) which was also found to be increased 70-fold in human iNeurons upon mitochondrial depolarisation (Ordureau et al., 2020).

Of the common substrates identified in our neuronal systems, a number have been previously quantified in HeLa and human iNeurons including CISD1, HK1, MFN1/2, RHOT1/2, TOMM70, and VDAC1/2/3 and their collective ubiquitylation together with other substrates has been associated with mitophagic signalling and clearance (Ordureau et al., 2020, Ordureau et al., 2018). Our analysis also found ATAD1 (Msp1) to be ubiquitylated upon Parkin activation (Figure S6). Akin to p97/Cdc48, Msp1 extracts misfolded proteins from the MOM providing a direct link between a Parkin substrate and mitochondrial quality control component (Wang and Walter, 2020, Wang et al., 2020). Interestingly we identified common ubiquitylation sites on FKBP8 (FK506 binding protein 8; also known as FKBP38) and SYNJ2BP (also known as OMP25) that are both anchored to the MOM via TOM-independent ‘tail anchor’ mitochondrial targeting signals within a C-terminal juxtamembrane sequence known as the CSS (Borgese and Fasana, 2011). The CSS of FKBP38 has also been demonstrated in MEFs to mediate its escape from the mitochondria to the ER during mitophagy thereby avoiding its degradation and this was also dependent on Parkin activity (Saita et al., 2013). However, in our studies we found that FKBP8 was regulated independent of PINK1 suggesting an alternate ubiquitin E3 ligase targets this protein and in future studies it will be interesting to assess whether FKBP8 undergoes escape to the ER in neurons and dissect the mechanism. Activation of Parkin in neurons may also be associated with proximal signalling events independent of mitophagy and interestingly, several of the common neuronal sites we identified, occur on proteins involved in fatty acid oxidation and metabolism in the mitochondria including CPT1α, ACSL1, VDACs, CYB5R3 (Figure 2 and 3). Recently several *in vivo* physiological studies have suggested a link between fatty acid oxidation and mitophagy mechanisms in non-neuronal tissues (Shao et al., 2020, Li et al., 2020) and in future work it will be interesting to investigate whether Parkin-induced ubiquitylation of these proteins affects fatty acid metabolism at the molecular level in neurons.

Since this is the first study to define common substrates regulated by human and mouse Parkin, we undertook a multiple sequence alignment analysis of the human Lys-targeted sequences to identify a putative recognition motif for Parkin, however this did not reveal a clear targeting sequence for Parkin-directed Lysine ubiquitylation (Figure S12) consistent with previous studies that have suggested that Parkin lacks sequence and structural specificity for substrate ubiquitylation (McWilliams et al., 2018, Ordureau et al., 2020, Ordureau et al., 2018, Sarraf et al., 2013).

Previous analysis of VDAC1 has suggested that the topology of the protein at the MOM and surface exposure of Lys residues to the cytosolic side strongly influences Parkin targeting however, it is not known if this is a general mechanism (Ordureau et al., 2020, Ordureau et al., 2018). Structural analysis to date has elaborated the mechanism of Parkin activation by PINK1 phosphorylation of Parkin and Ubiquitin (Gladkova et al., 2018, Sauve et al., 2018) and in future studies it will be interesting to reconstitute Parkin catalysed ubiquitylation of model substrates and solve structures of Parkin bound to E2 and substrate to uncover any autonomous structural determinants of Lysine recognition by Parkin and our elaboration of its key substrates in neurons will aid in these studies.

The ubiquitylation of MOM proteins on damaged mitochondria is followed by the recruitment of the autophagy receptor proteins NDP52, p62, OPTN, Tax1bp and NBR1 and several studies suggests that NDP52 and OPTN are the main ubiquitin-dependent receptors for efficient mitophagic clearance of damaged mitochondria both in HeLa cells and neurons (Heo et al., 2015, Evans and Holzbaur, 2020, Lazarou et al., 2015). The mechanism of how these are recruited to the MOM is not fully understood since NDP52 and OPTN have been demonstrated to bind diverse ubiquitin chain types including M1, K48 and K63 chains (Li et al., 2018, Xie et al., 2015) and there are conflicting reports on how recruitment of these receptors is affected by phospho-ubiquitin (Lazarou et al., 2015, Li et al., 2018, Ordureau et al., 2018, Richter et al., 2016). The UBAN domain of OPTN preferentially binds M1 or K63-linked Ub chains and previous analysis has found that phosphorylation of OPTN by TBK1 enhances its binding to Ub chains (Richter et al., 2016). Interestingly phosphorylation of Ser473 enhanced affinity of OPTN to longer K63 chains but also promoted binding to other linkage types which may not be relevant to mitophagic signalling in neurons (Richter et al., 2016), Heo et al., 2015). Our findings of defined substrates and specific accumulation of K63-linked ubiquitin chain types in neurons will be beneficial to studies underway to reconstitute mitophagy mechanisms in vitro (Shi et al., 2020a, Shi et al., 2020b) that may shed light on the initial steps of ubiquitin receptor recruitment to mitochondria. It was recently suggested that ubiquitin chains alone are sufficient to stimulate mitophagy (Yamano et al., 2020) and our findings also lend tractability to undertake genetic analysis as to whether ubiquitylation of specific substrates is critical for mitophagy or not. FKBP8 has also been reported to act as mitophagy receptor in HeLa cells via an N-terminal LC3-interacting region (LIR) that binds strongly to LC3A however this was independent of Parkin activation (Bhujabal et al., 2017) and future studies should be directed at its role in mitophagy in neurons in light of our discovery of Parkin dependent ubiquitylation.

A recent study has reported that two protein kinases, Cyclin G-associated kinase (GAK) and Protein Kinase C Delta (PRKCδ) are novel regulators of Parkin-independent mitophagy and recruit ULK1 to autophagosome structures close to mitochondria (Munson et al., 2020). Whilst we did not detect either GAK or PRKδ, we did detect multiple kinases in our mitochondrial enriched fractions that were ubiquitylated including PRKCβ, PRKCε and PRKC γ isoforms and in future work it will important to determine whether these and other kinases recruited to mitochondria regulate AO-induced mitophagy in neurons (Table S4). Our analysis has also highlighted PINK1-independent and Parkin-independent ubiquitylation sites following AO stimulation. Whilst NIX/BNIP3 and FUNDC1 receptor-mediated mitophagy have been linked to mitochondrial turnover during metabolic reprogramming or hypoxia, very little is known on PINK1-independent ubiquitin-mediated mechanisms regulating mitophagy (Montava-Garriga and Ganley, 2020). Our discovery of clear-cut PINK1-independent substrates suggest that there are parallel ubiquitin signalling pathways that are activated alongside PINK1-Parkin in neurons following AO treatment. We have compared our proteome dataset in neurons with the UbiHub online database (Liu et al., 2019) and found 40 E2, 165 ‘simple’ E3, and 155 ‘complex’ E3 components, that are present within our mitochondrial-enriched neuron fractions and we have deposited this data together with all detected ubiquitylation sites in a searchable online database that we have termed MitoNUb (https://shiny.compbio.dundee.ac.uk/MitoNUb/). Only two E3s, MUL1 and FBXL4, are located within the mitochondria and the remainder are cytosolic and in future work it will be interesting to define those that may be recruited and activated at the mitochondria upon AO treatment. Intriguingly DJ1 ubiquitylation was reduced by AO at Lys93 and to a lesser extent at Lys130 (Figure S5A). Both sites are highly conserved and lie close to Cys106 that is modified under oxidative stress conditions, thereby stimulating DJ1 recruitment from the cytosol to the mitochondria (Figure S13A) (Canet-Aviles et al., 2004). Patients with DJ1 mutations resemble PINK1 and Parkin patients (van der Vlag et al., 2020), however, we observed that the reduction of DJ1 ubiquitylation at Lys 93 was PINK1 independent (Figure S13B) and in future work it will be interesting to determine the mechanism of the functional impact of DJ1 ubiquitylation in the context of mitophagic signalling.

Overall this study together with previous findings in human iNeurons (Ordureau et al., 2020) provides a common signature of ubiquitylation events occurring in neuronal mitochondria that can be utilised to investigate Parkin activity in other neuronal cell types including dopaminergic neurons. The identification of a specific set of sites targeted by endogenous Parkin in cell types relevant to Parkinson’s will provide impetus to the development of facile proteomic and cell-based assays to monitor Parkin activity that will aid drug discovery and translational efforts to activate PINK1 and Parkin as a potential therapeutic strategy in Parkinson’s disease (Padmanabhan et al., 2019).

## STAR*METHODS

### KEY RESOURCES TABLE

#### Antibodies for biochemical studies

The following primary antibodies were used: Anti-Parkin phospho-Ser65 rabbit monoclonal antibody was raised by Epitomics/Abcam in collaboration with the Michael J Fox Foundation for Research (Please contact for questions). CISD1 (Cell signalling Technology, Proteintech), GAPDH (Santa Cruz), Ubiquitin (BioLegend), CPT1α (Abcam), HK1 (Thermo Fisher Scientific), GK (Abcam), DCAKD (Aviva Systems Biology), ABCD3 (Aviva Systems Biology), ACSL1 (Cell signalling Technology), ACSL6 (Sigma-Aldrich), AGPAT5 (Abcam), MARC2 Sigma-Aldrich), CYB5B (Novus Biologicals), CYB5R3 (Sigma), MFN1 (Abcam), MFN2 (Proteintech), RHOT2 (Proteintech), TOMM70 (Aviva Systems Biology), SLC25A46 (Proteintech), FAM213A (Novus Biologicals), MAO-A (Proteintech), MAO-B (Abcam), HSDL1 (Proteintech), CAMK2α (Thermo Fisher Scientific), CAMK2β (Thermo Fisher Scientific), DCAMKL2 (Abcam), CAD (Novus), PRKCγ (Proteintech), ATAD1/Thorase (NeuroMab), TDRKH (Proteintech), FBXO41 (Proteintech), SNX3 (Sigma-Aldrich), CNN3 (Sigma-Aldrich), SH3BP4 (Novus Biologicals), MAPRE2 (Proteintech), RAB5C (MyBiosource), p23 (Thermo Fisher Scientific), VPS35 (Abcam), OPA1 (Cell Signalling Technology), Rab8A (Cell Signallling Technology), Rab8A phospho-Ser111 (Abcam). Horseradish-peroxidase (HRP)-conjugated secondary antibodies (Sigma-Aldrich) were used.

#### Materials and Reagents

HaloLink resin was purchased from Promega. All mutagenesis was carried out using the QuikChange site-directed mutagenesis method (Stratagene) with KOD polymerase (Novagen). All DNA constructs were verified by MRC Protein Phosphorylation and Ubiquitylation Unit (PPU) DNA Sequencing Service, School of Life Sciences, University of Dundee, using DYEnamic ET terminator chemistry (Amersham Biosciences) on Applied Biosystems automated DNA sequencers. DNA for bacterial protein expression was transformed into *E. coli* BL21 DE3 RIL (codon plus) cells (Stratagene). Stock solutions of Antimycin A (Sigma), and Oligomycin (Sigma) were used for experiments in cells. Unless otherwise specified, general reagents and chemicals were from Sigma-Aldrich (Merck) and cell culture reagents were from Gibco/Invitrogen (Thermo Fisher Scientific). All cDNA plasmids, antibodies and recombinant proteins generated inhouse for this study are available on request through our dedicated reagents website: https://mrcppureagents.dundee.ac.uk/

### STAR METHOD DETAILS

#### Primary cortical neuron preparation and culture

Primary mouse cortical neurons were isolated from the brains of C57Bl/6J embryos of either sex at E16.5. Embryonic cortices were collected in HBSS, and cells were dissociated by incubation with trypsin-EDTA (#25300-054, Gibco) at 37°C. Cells were then diluted in Neurobasal medium containing B27 supplement, Glutamax, penicillin/streptomycin and plated at a density of 5.0 × 10^5^ cells/well on 6-well plates coated with 0.1 mg ml^−1^ poly-l-lysine (PLL; Sigma-Aldrich). Neurons were cultured at 37°C in a humidified incubator with 5% CO_2_. Every 5 days, the medium was replaced with fresh medium containing B27. To depolarize mitochondrial membrane potential in neurons, cultures were treated for 5 hours with 10 µM Antimycin A (Sigma-Aldrich) and 1 µM Oligomycin (Sigma-Aldrich) in DMSO at 37°C.

#### Isolation of mitochondrial enriched membrane fraction

Cells were collected in ice-cold PBS containing sodium orthovanadate (1 mM), sodium glycerophosphate (10 mM), sodium fluoride (50 mM), sodium pyrophosphate (10 mM), phenylmethylsulfonyl fluoride (PMSF; 0.1 mM) protease inhibitor cocktail, 200 mM chloroacetamide and centrifugated at 500g for 3 min at 4°C. Cell pellets were resuspended and lysed in homogenisation buffer containing 250 mM sucrose, 300 mM imidazole, 1mM sodium orthovanadate, phosphatase inhibitor cocktail 3 (Sigma-Aldrich), protease inhibitor cocktail (Roche) and supplemented with 200 mM chloroacetamide at 4°C. Cells were disrupted using a metal hand-held homogenizer (40 passes) and the lysates were clarified by centrifugation (2000 rpm at 4°C for 5 min). The supernatant was harvested and subjected to an additional centrifugation step at 40000 *rpm* for 1 hour at 4°C. The resulting pellet containing the membrane-enriched fraction was resuspended in Mitobuffer (270 mM sucrose, 20m M HEPES, 3 mM EDTA, 1 mM sodium orthovanadate, 10 mM sodium β-glycerophosphate, 50 mM NaF, 5 mM sodium pyrophosphate, pH 7.5 and protease inhibitor cocktail (Roche) supplemented with 200 mM chloroacetamide at 4°C) and solubilised with a probe sonicator (5 seconds, 20% Amplitude).

#### Whole cell lysate preparation

Primary cortical neurons and SH-SY5Y cells were sonicated in lysis buffer containing Tris⋅HCl (50 mm, pH 7.5), EDTA (1 mM), ethylene glycol bis(β-aminoethyl ether)-*N*,*N*,*N*′,*N*′-tetraacetic acid (EGTA; 1 mM), Triton (1[%, w/v), sodium orthovanadate (1 mM), sodium glycerophosphate (10 mM), sodium fluoride (50 mM), sodium pyrophosphate (10 mM), sucrose (0.25□mM), benzamidine (1 mM), phenylmethylsulfonyl fluoride (PMSF; 0.1 mM) protease inhibitor cocktail (Roche), phosphatase inhibitor cocktail 2 and 3 (Sigma-Aldrich) and Chloracetamide (200 mM). Following sonication, lysates were incubated for 30 min on ice. Samples were spun at 20□800□*g* in an Eppendorf 5417R centrifuge for 30 min. Supernatants were collected and protein concentration was determined by using the Bradford kit (Pierce).

#### Ubiquitin enrichment

For ubiquitylated protein capture, Halo□tagged ubiquitin□binding domains (UBDs) of TUBE (tandem□repeated ubiquitin□binding entities), multi-DSK (yeast Dsk1 ubiquitin binding protein) and multi-DSK mutant were incubated with HaloLink resin (200 μL, Promega) in binding buffer (50 mm Tris⋅HCl, pH 7.5, 150 mm NaCl, 0.05□% NP□40) overnight at 4□°C. Membrane-enriched fraction (400 μg) was used for pulldown with HALO□UBDs. Halo Tube beads (20 μL) were added to neuronal membrane enrichment and incubated for ON at 4□°C. The beads were washed three times with lysis buffer containing 0.25_m NaCl and eluted by resuspension in 1×LDS sample buffer (20 μL) with 1 mM dithiothreitol (DTT) or 2.5% 2-mercaptoethanol.

#### PINK1 immunoprecipitation

For immunoprecipitation of endogenous PINK1, 500 µg of whole-cell lysate or membrane fraction were incubated overnight at 4°C with 10 µg of PINK1 antibody (S774C, MRC PPU reagents and Services) coupled to Protein A/G beads (10 µl of beads per reaction, Amintra). The immunoprecipitants were washed three times with lysis buffer containing 150 mM NaCl and eluted by resuspending in 10 µl of 2× LDS sample buffer and incubating at 37°C for 15 min under constant shaking (2000 rpm) followed by the addition of 2.5% (by vol) 2-mercaptoethanol.

#### Immunoblotting

All the samples were subjected to SDS□PAGE (Bis-Tris 4–12□% gels), CAD and Nav1 proteins were separated by using Tris-Acetate (3-8% gels) and all the gels transferred onto Protran 0.45 PVDF membranes (Immobilon-P). Membranes were blocked for 1 h at room temperature with 5□% non□fat milk or bovine serum albumin (BSA) in Tris□buffered saline (TBST; 50 mm Tris·HCl and 150 mm NaCl, pH 7.5) containing 0.1□% Tween□20 and probed with the indicated antibodies overnight at 4□°C. Detection was performed using horseradish peroxidase (HRP)□conjugated secondary antibodies and enhanced chemiluminescence reagent.

#### Ubiquitylation Assay

In vitro ubiquitylation assays were performed using recombinant proteins purified from *E. coli* unless stated otherwise. 0.75 μM of wild type Parkin was incubated for 30 minutes at 37 °C in a thermo shaker at 1000 rpm with 0.36 μM wild type or kinase inactive (D359A) MBP-TcPINK1 (*Tribolium castaneum* PINK1) in 15 μl of kinase buffer [50 mM Tris-HCl pH 7.5, 0.1 mM EGTA, 10 mM MgCl_2_, and 0.1 mM ATP]. The ubiquitin master mix [50 mM Tris-HCl, 10 mM MgCl_2_, 2 mM ATP, 0.12 μM His-UbE1 expressed in Sf21 insect cells, 1 μM human UbE2L3, 50 μM Flag-ubiquitin, and 0.5 μM substrate] was added to a final volume of 30 μl and the reaction was incubated at 37 °C for 30 min in a thermo shaker at 1000 rpm. Reactions were terminated by the addition of 4 x LDS loading buffer and 5 μl of the final volume was resolved using SDS-PAGE on a 4-12 % Bis-tris gel and immunoblotted using an anti-FLAG, anti-His, or antibodies against the substrate being tested.

#### Immunoblotting of ubiquitylation assay components

Samples were resolved using SDS-PAGE on 4 to 12 % Bis-Tris gels in MOPS buffer and transferred to nitrocellulose membranes. Membranes were blocked with 5 % w/v milk powder in TBS + 0.1 % TWEEN 20 (TBS-T) for 1 h at room temperature then immunoblotted against the primary antibody in 5 % w/v BSA / TBS-T at 5 to 7 °C overnight. Protein bands were detected by blotting against secondary antibodies labelled with 800 nm or 680 nm fluorophores in TBS-T for 1 h at room temperature and imaged using LiCor.

#### Recombinant Protein Expression

##### Parkin

His_6_-SUMO cleaved wild type Parkin was expressed based upon the method (Kazlauskaite et al., 2014b). Briefly, plasmids were transformed in BL21 Codon Plus (DE3)-RIL *E. coli*, overnight cultures were prepared and used to inoculate 12 x 1 L of LB medium containing 50 μg/ml carbenicillin and 0.25 mM ZnCl_2_. These were initially incubated at 37 °C until the cultures reached an OD_600_ of 0.4, the incubator temperature was lowered to 15 °C, and once cultures reached an OD_600_ of 0.8-0.9 expression was induced by the addition of 25 μM IPTG. After overnight incubation (16 h) cells were pelleted by centrifugation (4200 x g, 25 minutes), the media was removed, and the cell pellet was suspended in lysis buffer [50 mM Tris-HCl pH 7.5, 250 mM NaCl, 15 mM imidazole (pH7.5), and 0.1 mM EDTA with 1 μM AEBSF and 10 μg/ml leupeptin added]. Cells were burst by sonication and cell debris were pelleted by centrifugation (35000 x g for 30 minutes at 4°C) and the supernatant was incubated with Ni-NTA resin for 1 hour at 5-7°C. Ni-NTA resin was washed 5 times in 7 x the resin volume of lysis buffer, and twice in 7 x the resin volume of cleavage buffer [50 mM Tris pH 8.3, 200 mM NaCl, 10 % glycerol and 1 mM TCEP]. Parkin was cleaved from the resin at 4°C overnight by the addition of a 1:5 mass ratio of His-SENP1 to total protein mass bound to the resin. After cleavage Parkin was concentrated and further purified using size exclusion chromatography on a Superdex S200 column (16/600). Parkin was eluted after 80 to 90 ml, fractions were pooled and concentrated the purity was tested using SDS-PAGE.

##### Cpt1α

A catalytic domain-containing fragment of Cpt1α (residues 145-773; missing the transmembrane domain) was expressed as His-SUMO tagged fusion protein. The cDNA of the protein coding sequences was subcloned into pET15b plasmid vectors and transformed into BL21-DE3 *E. Coli*. A 150 ml starter culture was inoculated using a swipe from the cell plate and incubated at 37 °C overnight. 12 x 1L LB media containing 50 μg/mL of carbenicillin were each inoculated with 20 ml of started culture. When cell cultures reached an OD600 of 0.8-0.9 and a temperature of 15 °C expression was induced by the addition of 50 μM IPTG. Cell cultures were left to express at 15 °C overnight (∼16 h) then harvested by centrifugation at 5020 x g for 25 minutes; the media was removed, and the cell pellet was suspended in 25 ml per 1 L (∼5mL pellet) of lysis buffer [50 mM Tris-HCl pH 7.5, 250 mM NaCl, 15 mM imidazole (pH7.5), and 0.1 mM EDTA with 1 μM AEBSF and 10 μg/mL leupeptin added]. Cells were burst by sonication at 50 % amplitude on ice using 6 x 15 s pulses and 10 % glycerol was added and the lysis buffer was made up to 500 mM NaCl. Cell debris were pelleted by centrifugation at 35000 x g and 4 °C for 30 min, then the supernatant layer was transferred to Ni-NTA resin. The supernatant was incubated with Ni-NTA resin for 1 hour at 5-7 °C, washed 6 times in 7 x the resin volume of wash buffer [50 mM Tris-HCl pH 7.5, 500 mM NaCl, 15 mM imidazole (pH7.5), 0.5 mM TCEP, and 10 % glycerol] then His-SUMO-Cpt1α 145-773 was eluted by incubation with 4 x the resin volume of elution buffer [wash buffer containing 400 mM Imidazole] at 5-7 °C for 1 h. The solvent layer was dialysed against 5 L dialysis buffer [50 mM Tris-HCl pH 7.5, 200 mM NaCl, 0.5 mM TCEP, and 10 % glycerol] overnight at 4 °C with a 1 : 10 mass ratio of His-SENP1 : eluted protein mass added. Uncleaved Cpt1α and contaminants were depleted by incubating with Ni-NTA resin for 1 h at 5-7 °C. The solvent layer was concentrated and purified using size exclusion chromatography in dialysis buffer using a 16/600 Superdex SD200 column. Fractions containing Cpt1α 145-773 (peak at ∼75 ml) were pooled and concentrated to give the recombinant.

##### GST-Miro1

GST-Miro1 cDNA was subcloned into a pGEX6 plasmid vector and transformed into BL21 Codon Plus (DE3)-RIL *E. coli*. A 150 mL starter culture was inoculated using a swipe from the cell plate and incubated at 37 °C overnight. 12 x 1L LB media containing 50 μg /mL of carbenicillin were each inoculated with 20 mL of started culture. When cell cultures reached an OD600 of 0.8-0.9 and a temperature of 15°C expression was induced by the addition of 50 μM IPTG. Cell cultures were left to express at 15 °C overnight (∼16h) then harvested by centrifugation at 5020 x g for 25 min, the media was removed, and the cell pellet was suspended in 25 mL per 1 L (∼5mL pellet) of lysis buffer [50 mM Tris-HCl pH 7.5, 250 mM NaCl, 1 mM β-mercaptoethanol and 0.1 mM EDTA with 1μM AEBSF and 10 μg/mL leupeptin added]. Cells were burst by sonication at 50 % amplitude on ice using 6 x 15 s pulses. Cell debris were pelleted by centrifugation at 35000 x g and 4°C for 30 min, then the supernatant layer was transferred to glutathione resin. The supernatant was incubated with the GSH resin for 1 h at 5-7 °C, the GSH resin was washed 6 times in 14 x the resin volume of wash buffer [50 mM Tris-HCl pH 7.5, 200 mM NaCl, 0.5 mM TCEP, and 10 % glycerol], and the protein was eluted by incubation with wash buffer containing 10 mM glutathione for 1 h at 5-7 °C. Eluted supernatant was dialysed against 5L wash buffer at 5-7 °C overnight, concentrated and the final sample was flash frozen.

##### ACSL1 and SNX3

Cleaved ACSL1 (missing the N-terminal transmembrane domain) and SNX3, were expressed as a His_6_-SUMO tagged constructs. cDNA of the protein coding sequences were subcloned into pET15b plasmid vectors and transformed into BL21 Codon Plus (DE3)-RIL E*. coli*. A 100 mL starter culture was inoculated using a swipe from the cell plate and incubated at 37 °C overnight. 6 x 1L LB media containing 50 μg/mL of carbenicillin were each inoculated with 20 mL of started culture. When cell cultures reached an OD600 of 0.8-0.9 and a temperature of 15°C expression was induced by the addition of 100 μM IPTG. Cell cultures were left to express at 15 °C overnight (∼16h) then harvested by centrifugation at 5020 x g for 25 minutes, the media was removed, and the cell pellet was suspended in 25 mL per 1 L (∼5mL pellet) of lysis buffer [50 mM Tris-HCl pH 7.5, 250 mM NaCl, 15 mM imidazole (pH7.5), and 0.1 mM EDTA with 1 μM AEBSF and 10 μg/mL leupeptin added]. Cells were burst by sonication at 50 % amplitude on ice using 6 x 15 s pulses and 10 % glycerol was added and the lysis buffer was made up to 500 mM NaCl. Cell debris were pelleted by centrifugation at 35000 x g and 4 °C for 30 min, then the supernatant layer was transferred to Ni-NTA resin. The supernatant was incubated with Ni-NTA resin for 1 h at 5 −7 °C, washed 6 times in 7 x the resin volume of wash buffer [50 mM Tris-HCl pH 7.5, 500 mM NaCl, 15 mM imidazole (pH7.5), 0.5 mM TCEP, and 10 % glycerol] then 2 times in 7 x the resin volume of cleavage buffer [50 mM Tris-HCl pH 7.5, 200 mM NaCl, 0.5 mM TCEP, and 10 % glycerol]. The resin was incubated overnight without agitation at 4°C in 4 x the resin volume of cleavage buffer with a 1 : 10 mass ratio of His-Senp1 : bound protein. The solvent layer was depleted against a further 1 mL Ni-NTA resin at 5-7°C for 1 h. The final solution was concentrated, flash frozen in liquid nitrogen, and stored at −80 °C.

##### MBP-CamK2α and MBP-CamK2

CamK2α and CamK2β were expressed as MBP tagged constructs. cDNA of the protein coding sequences were subcloned into pMEX3Cb plasmid vectors and transformed into BL21 Codon Plus (DE3)-RIL *E. coli*. A 100 mL starter culture was inoculated using a swipe from the cell plate and incubated at 37°C overnight. 6 x 1L LB media containing 50 μg/mL of carbenicillin were each inoculated with 20 mL of started culture. When cell cultures reached an OD600 of 0.8-0.9 and a temperature of 15°C expression was induced by the addition of 400 μM IPTG. Cell cultures were left to express at 15°C overnight (∼16h) then harvested by centrifugation at 5020 x g for 25 min, the media was removed, and the cell pellet was suspended in 25 mL per 1 L (∼5mL pellet) of lysis buffer [50 mM Tris-HCl pH 7.5, 250 mM NaCl, and 0.1 mM EDTA with 1 μM AEBSF and 10 μg/mL leupeptin added]. Cells were burst by sonication at 50 % amplitude on ice using 6 x 15 s pulses and 10 % glycerol was added and the lysis buffer was made up to 500 mM NaCl. Cell debris were pelleted by centrifugation at 35000 x g and 4°C for 30 min. The supernatant was incubated with amylose resin for 1 h at 5-7 °C, washed 6 times in 7 x the resin volume of lysis buffer then eluted in elution buffer [50 mM Tris-HCl pH 7.5, 500 mM NaCl, 10 mM Maltose, 0.5 mM TCEP, and 10 % glycerol]. The eluted sample was dialysed overnight against 5L of sample buffer [50mM Tris-HCl pH 7.5, 0.1mM EGTA, 150mM NaCl, 0.1% ß-Mercaptoethanol, 270mM sucrose, 0.03% Brij-35]. The final solution was concentrated, flash frozen in liquid nitrogen, and stored at −80°C.

##### GST-Fam213A, GST-MAOA and GST-MAOB

Fam213A, MAOA and MAOB were expressed as GST tagged constructs. cDNA of the protein coding sequences were subcloned into pGEX6 plasmid vectors and transformed into BL21 Codon Plus (DE3)-RIL Ecoli. A 100 mL starter culture was inoculated using a swipe from the cell plate and incubated at 37°C overnight. 6 x 1L LB media containing 50 μg/mL of carbenicillin were each inoculated with 20 mL of started culture. When cell cultures reached an OD600 of 0.8-0.9 and a temperature of 15 °C expression was induced by the addition of 50 μM IPTG. Cell cultures were left to express at 15 °C overnight (∼16h) then harvested by centrifugation at 5020 x g for 25 min, the media was removed, and the cell pellet was suspended in 25 mL per 1 L (∼5mL pellet) of lysis buffer [50 mM Tris-HCl pH 7.5, 250 mM NaCl, and 0.1 mM EDTA with 1 μM AEBSF and 10 μg/mL leupeptin added]. Cells were burst by sonication at 50 % amplitude on ice using 6 x 15 s pulses and 10 % glycerol was added and the lysis buffer was made up to 500 mM NaCl. Cell debris were pelleted by centrifugation at 35000 x g and 4 °C for 30 min. The supernatant was incubated with GSH resin for 1 h at 5-7 °C, washed 6 times in 7 x the resin volume of lysis buffer then eluted in elution buffer [50 mM Tris-HCl pH 7.5, 500 mM NaCl, 10 mM reduced-glutathione, 0.5 mM TCEP, and 10 % glycerol]. The eluted sample was dialysed overnight against 5L of sample buffer [50mM Tris-HCl pH 7.5, 0.1mM EGTA, 150mM NaCl, 0.1% β-Mercaptoethanol, 270mM sucrose, 0.03% Brij-35]. The final solution was concentrated, flash frozen in liquid nitrogen, and stored at −80 °C.

#### Online Database

diGLY peptide TMT ratios were normalised to the total proteome. For differential expression, these data were normalised to median in each sample, to account for sample-to-sample variation. Differential expression was performed with *limma* (Ritchie et al., 2015) using logarithms of normalised peptide ratios. P-values were corrected for multiple tests using Benjamini-Hochberg method. All analysis was done in R and the code is available at https://github.com/bartongroup/MG_UbiMito.

An online interactive tool was created using Shiny framework and it is publicly available at https://shiny.compbio.dundee.ac.uk/MitoNUb/. It allows for selection of peptides based on differential expression significance, fold change, presence or absence in Mitocarta and/or total proteome. Selected peptides provide instantaneous GO-term and Reactome pathway enrichment results.

#### Proteomics – General Sample Preparation

Protein extracts lysed in 8M urea, were subjected to disulfide bond reduction with 5 mM TCEP (room temperature, 10 min) and alkylation with 25 mM chloroacetamide (room temperature, 20 min). Methanol–chloroform precipitation was performed prior to protease digestion. In brief, four parts of neat methanol were added to each sample and vortexed, one part chloroform was then added to the sample and vortexed, and finally three parts water was added to the sample and vortexed. The sample was centrifuged at 6 000 rpm for 2 min at room temperature and subsequently washed twice with 100% methanol. Samples were resuspended in 100 mM EPPS pH8.5 containing 0.1% RapiGest and digested at 37°C for 4h with LysC protease at a 200:1 protein-to-protease ratio. Trypsin was then added at a 100:1 protein-to-protease ratio and the reaction was incubated for a further 6 h at 37°C. Samples were acidified with 1% Formic Acid for 15 min and subjected to C18 solid-phase extraction (SPE) (Sep-Pak, Waters). The Pierce Quantitative Colorimetric Peptide Assay (cat.no. 23275) was used to quantify the digest and to accurately aliquot the desired amount of peptides per sample needed for downstream application.

#### Proteomics – Total proteomics analysis using TMT

Tandem mass tag labeling of each sample (100 μg peptide input) was performed by adding 10 μL of the 20 ng/μL stock of TMT reagent along with acetonitrile to achieve a final acetonitrile concentration of approximately 30% (v/v). Following incubation at room temperature for 1 h, the reaction was quenched with hydroxylamine to a final concentration of 0.5% (v/v) for 15 min. The TMT-labeled samples were pooled together at a 1:1 ratio. The sample was vacuum centrifuged to near dryness, and subjected to C18 solid-phase extraction (SPE) (50 mg, Sep-Pak, Waters).

Dried TMT-labeled sample was resuspended in 100 μL of 10 mM NH_4_HCO_3_ pH 8.0 and fractionated using BPRP HPLC (Wang et al., 2011). Briefly, samples were offline fractionated over a 90 min run, into 96 fractions by high pH reverse-phase HPLC (Agilent LC1260) through an aeris peptide xb-c18 column (Phenomenex; 250 mm x 3.6 mm) with mobile phase A containing 5% acetonitrile and 10 mM NH_4_HCO_3_ in LC-MS grade H_2_O, and mobile phase B containing 90% acetonitrile and 10 mM NH_4_HCO_3_ in LC-MS grade H_2_O (both pH 8.0). The 96 resulting fractions were then pooled in a non-continuous manner into 24 fractions (as outlined in Figure S5 of (Paulo et al., 2016a)) and 12 fractions (even numbers) were used for subsequent mass spectrometry analysis. Fractions were vacuum centrifuged to near dryness. Each consolidated fraction was desalted via StageTip, dried again via vacuum centrifugation, and reconstituted in 5% acetonitrile, 1% formic acid for LC-MS/MS processing.

Mass spectrometry data were collected using an Orbitrap Fusion Lumos mass spectrometer (Thermo Fisher Scientific, San Jose, CA) coupled to a Proxeon EASY-nLC1200 liquid chromatography (LC) pump (Thermo Fisher Scientific). Peptides were separated on a 100 μ inner diameter microcapillary column packed in house with ∼35 cm of Accucore150 resin (2.6 μm, 150 Å, ThermoFisher Scientific, San Jose, CA) with a gradient consisting of 5%–21% (0-125 min), 21%–28% (125-140min) (ACN, 0.1% FA) over a total 150 min run at ∼500 nL/min. For analysis, we loaded 1/10 of each fraction onto the column. Each analysis used the Multi-Notch MS^3^-based TMT method (McAlister et al., 2014), to reduce ion interference compared to MS^2^ quantification (Paulo et al., 2016b). The scan sequence began with an MS^1^ spectrum (Orbitrap analysis; resolution 120,000 at 200 Th; mass range 400−1400 m/z; automatic gain control (AGC) target 5 × 10^5^; maximum injection time 50 ms). Precursors for MS^2^ analysis were selected using a Top10 method. MS^2^ analysis consisted of collision-induced dissociation (quadrupole ion trap analysis; Turbo scan rate; AGC 2.0 × 10^4^; isolation window 0.7 Th; normalized collision energy (NCE) 35; maximum injection time 90 ms). Monoisotopic peak assignment was used and previously interrogated precursors were excluded using a dynamic window (150 s ± 7 ppm) and dependent scans were performed on a single charge state per precursor. Following acquisition of each MS^2^ spectrum, a synchronous-precursor-selection (SPS) MS^3^ scan was collected on the top 10 most intense ions in the MS^2^ spectrum (McAlister et al., 2014). MS^3^ precursors were fragmented by high energy collision-induced dissociation (HCD) and analyzed using the Orbitrap (NCE 65; AGC 3 × 10^5^; maximum injection time 150 ms, resolution was 50,000 at 200 Th).

#### Immunoprecipitation of diGLY-Containing Peptides

diGLY capture was performed largely as described (<u>Rose et al., 2016</u>). The diGly monoclonal antibody (Cell Signaling Technology; D4A7 clone) (32 μg antibody/1 mg peptide) was coupled to Protein A Plus Ultralink resin (1:1 μL slurry/ μg antibody) (Thermo Fisher Scientific) overnight at 4°C prior to its chemical cross-linking reaction. Dried peptides (1 mg starting material) were resuspended in 1.5 mL of ice-cold IAP buffer [50 mM MOPS (pH 7.2), 10 mM sodium phosphate and 50 mM NaCl] and centrifuged at maximum speed for 5 min at 4°C to remove any insoluble material. Supernatants (pH ∼7.2) were incubated with the antibody beads for 2 hr at 4°C with gentle end-over-end rotation. After centrifugation at 215 × g for 2 min, beads were washed three more times with ice-cold IAP buffer and twice with ice-cold PBS. The diGLY peptides were eluted twice with 0.15% TFA, desalted using homemade StageTips and dried via vacuum centrifugation, prior to TMT labeling.

#### Proteomics – diGLY proteomics analysis using TMT

TMT-labeled diGLY peptides were fractionated according to manufacturer’s instructions using High pH reversed-phase peptide fractionation kit (Thermo Fisher Scientific) for a final 6 fractions and subjected to C18 StageTip desalting prior to MS analysis.

Mass spectrometry data were collected using an Orbitrap Fusion Lumos mass spectrometer (Thermo Fisher Scientific, San Jose, CA) coupled to a Proxeon EASY-nLC1200 liquid chromatography (LC) pump (Thermo Fisher Scientific). Peptides were separated on a 100 μm inner diameter microcapillary column packed in house with ∼35 cm of Accucore150 resin (2.6 μm, 150 Å ThermoFisher Scientific, San Jose, CA) with a gradient consisting of 3%–26% (0-130 min), 26%–32% (130-140min) (ACN, 0.1% FA) over a total 150 min run at ∼500 nL/min. For analysis, we loaded 1/2 of each fraction onto the column. Each analysis used the Multi-Notch MS^3^-based TMT method (McAlister et al., 2014). The scan sequence began with an MS^1^ spectrum (Orbitrap analysis; resolution 120,000 at 200 Th; mass range 400−1250 m/z; automatic gain control (AGC) target 1 × 10^6^; maximum injection time 100 ms). Precursors for MS^2^ analysis were selected using a Top 4 s method. MS^2^ analysis consisted of collision-induced dissociation (quadrupole Orbitrap analysis; AGC 1 × 10^5^; isolation window 0.7 Th; normalized collision energy (NCE) 35; maximum injection time 300 ms resolution was 7,500 at 200 Th). Monoisotopic peak assignment was used and previously interrogated precursors were excluded using a dynamic window (120 s ± 7 ppm). As described previously, only precursors with a charge state between 3 and 6 were selected for downstream analysis (Rose et al., 2016). Following acquisition of each MS^2^ spectrum, a synchronous-precursor-selection (SPS) MS^3^ scan was collected on the top 10 most intense ions in the MS^2^ spectrum (McAlister et al., 2014). MS^3^ precursors were fragmented by high energy collision-induced dissociation (HCD) and analyzed using the Orbitrap (NCE 65; AGC 2 × 10^5^; maximum injection time 500 ms, resolution was 50,000 at 200 Th).

#### Proteomics – Data analysis

Mass spectra were processed using a Sequest-based or Comet-based (2014.02 rev. 2) in-house software pipeline (Eng et al., 2013, Huttlin et al., 2010). Spectra were converted to mzXML using a modified version of ReAdW.exe. Database searching included all entries from the Mouse Reference Proteome (2017-05) UniProt database, as well as an in-house curated list of contaminants. This database was concatenated with one composed of all protein sequences in the reversed order. Searches were performed using a 20 ppm precursor ion tolerance for total protein level analysis. The product ion tolerance was set to 0.9 Da (0.03 Da for diGLY searches). These wide mass tolerance windows were chosen to maximize sensitivity in conjunction with Sequest searches and linear discriminant analysis (Beausoleil et al., 2006, Huttlin et al., 2010). TMT tags on lysine residues and peptide N termini (+229.163 Da) and carbamidomethylation of cysteine residues (+57.021 Da) were set as static modifications, while oxidation of methionine residues (+15.995 Da) was set as a variable modification. For phosphorylation dataset search, phosphorylation (+79.966 Da) on Serine or Threonine and deamidation (+0.984 Da) on Asparagine or Glutamine were set as additional variable modifications. For diGLY dataset search, GlyGly modification (+114.0429 Da) was also set as a variable modification. Peptide-spectrum matches (PSMs) were adjusted to a 1% false discovery rate (FDR) (Elias and Gygi, 2007). PSM filtering was performed using a linear discriminant analysis, as described previously (Huttlin et al., 2010), while considering the following parameters: XCorr (or Comet Log Expect), ΔCn (or Diff Seq. Delta Log Expect), missed cleavages, peptide length, charge state, and precursor mass accuracy. For TMT-based reporter ion quantitation, we extracted the summed signal-to-noise (S:N) ratio for each TMT channel and found the closest matching centroid to the expected mass of the TMT reporter ion (integration tolerance of 0.003 Da). For protein-level comparisons, PSMs were identified, quantified, and collapsed to a 1% peptide false discovery rate (FDR) and then collapsed further to a final protein-level FDR of 1%. Moreover, protein assembly was guided by principles of parsimony to produce the smallest set of proteins necessary to account for all observed peptides. Phosphorylation or ubiquitylation site localization was determined using the AScore algorithm (Beausoleil et al., 2006). AScore is a probability-based approach for high-throughput protein phosphorylation site localization. Specifically, a threshold of 13 corresponded to 95% confidence in site localization. Proteins and phosphorylated or ubiquitylated peptides were quantified by summing reporter ion counts across all matching PSMs using in-house software, as described previously (Huttlin et al., 2010). PSMs with poor quality, MS^3^spectra with isolation specificity less than 0.7, or with TMT reporter summed signal-to-noise ratio that were less than 150, or had no MS^3^ spectra were excluded from quantification (McAlister et al., 2012).

Protein or peptide quantification values were exported for further analysis in Microsoft Excel, GraphPad Prism and Perseus (Tyanova et al., 2016). For whole proteome analysis, each reporter ion channel was summed across all quantified proteins and normalized assuming equal protein loading of all samples. For diGly samples, the data was normalized to each individual protein abundance measured in parallel when available to correct for variation in protein abundance between treatments. Supplemental data Tables list all quantified proteins as well as associated TMT reporter ratio to control channels used for quantitative analysis.

Annotations for bona fide organellar protein markers were assembled using the proteins which had scored with confidence “very high” or “high” from the HeLa dataset previously published Itzhak D.N. (Itzhak et al., 2016). The following database containing mitochondrial protein were used: MitoCarta 3.0 (Rath et al., 2021).

#### Mitochondrial UB and Poly-UB Capture and Proteomics

Mitochondrially derived ubiquitylated proteins were purified using Halo-4×UBA^UBQLN1^ and Halo-5xUBA^DSK2^ as described (Ordureau et al., 2015, Ordureau et al., 2014). Briefly, whole-cell extracts (0.5 mg) or mitochondrial extracts (0.5 mg) that were lysed in lysis buffer containing 50 mM chloroacetamide were incubated at 4°C for 16h with 25 μL of Halo-4×UBA^UBQLN1^ beads (pack volume). Subsequently, the supernatant was incubated with 15 μ of Halo-5xUBA^DSK2^ for 2 hours to capture any possible residual mono-ubiquitylated proteins. Halo beads were combined and following 4 washes with lysis buffer containing 0.5 M NaCl and one final wash in 10 mM Tris pH 8.0, proteins were released from the Halo-UBA resin using 6 M guanidine HCL. Samples were subjected to TCA precipitation and digested overnight at 37 °C with Lys-C and trypsin [in 100 mM tetraethylammonium bromide, 0.1% Rapigest (Waters Corporation), 10% (vol/vol) acetonitrile (ACN)]. Digests were acidified with an equal volume of 5% (vol/vol) formic acid (FA) to a pH of ∼2 for 30 min, dried down, resuspended in 5% (vol/vol) FA, and subjected to C18 StageTip (packed with Empore C18; 3M Corporation) desalting. Samples were analyzed by liquid chromatography (LC)/tandem MS for AQUA proteomics as described below.

#### UB-AQUA Proteomics

UB-AQUA was performed largely as described previously but with several modifications (Ordureau et al., 2015, Ordureau et al., 2014). A collection of 21 heavy-labeled reference peptides (Ordureau et al., 2015, Ordureau et al., 2014), each containing a single ^13^C/^15^N-labeled amino acid, was produced at Cell Signaling Technologies and quantified by amino acid analysis. UB-AQUA peptides from working stocks [in 5% (vol/vol) FA] were diluted into the digested sample [in 5% (vol/vol) FA] to be analyzed to an optimal final concentration predetermined for in-dividual peptide such that each peptide’s intensity would range between 10^6^ and 10^8^. Samples and AQUA peptides were oxidized with 0.1% hydrogen peroxide for 30 min, subjected to C18 StageTip and resuspended in 5% (vol/vol) FA. Replicate experiments were performed and analyzed sequentially by LC/MS on an Orbitrap Fusion Lumos instrument coupled to an Easy-nLC 1200 (Thermo Fisher Scientific) ultra-high-pressure liquid chromatography (UHPLC) pump. Peptides were separated on a 100 μm inner diameter microcapillary column packed in house with ∼35 cm of Accucore150 resin (2.6 μm, 150 Å, ThermoFisher Scientific, San Jose, CA). The column was equilibrated with buffer A (3% ACN + 0.125% FA). Peptides were loaded onto the column in 100% buffer A. Separation and elution from the column were achieved using a 75-min 0–28% gradient of buffer B [100% (vol/vol) ACN + 0.125% FA]. The scan sequence began with FTMS^1^ spectra (resolution of 120,000; mass range 300-1000 m/z; automatic gain control (AGC) target 5×10^5^, max injection time of 100 ms). The second scan sequence consisted of a targeted-MS^2^ (tMS^2^) method were MS^2^ precursors of interest were isolated using the quadrupole and analyzed in the Orbitrap (FTMS^2^) with a 0.7 Th isolation window, 30k resolution, 5×10^4^ AGC target and a max injection time of 54 ms. MS2 precursors were fragmented by HCD at a normalized collision energy (NCE) of 32%. LC-MS data analysis was performed using Skyline software (MacLean et al., 2010) with manual validation of precursors and fragments. The results exported to Excel and GraphPad Prism for further analysis and plotting. Total UB was determined as the average of the total UB calculated for each individual locus, unless specified otherwise.

#### Proteomics – Ubiquitylated site ranking analysis

In addition to TMT-based reporter ion quantitation, we also extracted the MS1 precursor abundance (intensity-based) for each diGLY peptide, a value indicative of the relative abundance of the peptide in the tryptic sample. Each MS1-based abundance measured should be a representation of the sum of all the respective TMT-labeled peptides combined. Therefore, for a rudimentary metric of site abundance across samples, we divided the total MS1-abundance for individual diGLY peptides by their respective TMT summed signal-to-noise ratio to each TMT channel.

#### Mouse breeding and maintenance

The C57BL/6J mice were obtained from Charles River Laboratories (Kent-UK), the *PINK1* and *Parkin* knockout mouse models used in this study were generated as previously described (McWilliams et al., 2018) housed in a SPF facility in temperature-controlled rooms at 21°C, with 45-65% relative humidity and 12-h light/dark cycles. Mice had ad libitum access to food and water and regularly monitored by the School of Life Science Animal Unit Staff. All animal studies and breeding in Dundee was approved by the University of Dundee ethical review committee, and further subjected to approved study plans by the Named Veterinary Surgeon and Compliance Officer (Dr. Ngaire Dennison) and performed under a UK Home Office project license in accordance with the Animal Scientific Procedures Act (ASPA, 1986).

---

**Figure S1. Characterisation of PINK1 signalling in mouse cortical neurons.**

**A.**Time course analysis of phospho-Ser65 ubiquitin levels in C56BL/6J mouse cortical neurons upon AO stimulation. Membrane lysate were subjected to Halo-multiDSK (mDSK) pull-down assay and immunoblotting with anti-phospho-Ser65 ubiquitin and anti-ubiquitin antibodies. PINK1 stabilisation detected by immunoprecipitation-immunoblot. Immunoblot analysis with anti-Parkin, anti-phospho-Ser65 Parkin, anti-phospho-Ser111 RAB8A, anti-RAB8A and anti-GAPDH antibodies.

**B.** Comparison of phospho-Ser65 ubiquitin detection using Halo-TUBE and/or Halo-multiDSK (mDSK) proteins enrichment. Mutant non-binding form of Halo-multiDSK (mDSK) used as negative control for the ubiquitin pull-down assay. Immunoblotting with anti-ubiquitin antibody was used as loading control.

**C.** Immunoblots showing comparative analysis of phospho-Ser65 Ubiquitin levels in primary cortical neuron cultures from wild-type and Parkin knockout (KO) mice. Membrane lysate were enriched for ubiquitin by incubating with Halo-multiDSK (mDSK). Enriched lysates were subjected to immunoblotting with anti-phospho-Ser65 ubiquitin and anti-ubiquitin antibodies. IP-immunoblot showed PINK1 protein stabilisation after mitochondrial depolarisation. Membrane lysate were also subjected to SDS-PAGE and immunoblot analysis with anti-Parkin, anti-phospho-Ser65 Parkin, anti-phospho-Ser111 RAB8A, anti-RAB8A and anti-GAPDH antibodies. * Non-specific band.

**Figure S2. Parkin activation is not affected in VPS35 D620N neurons.**

Phospho-Ser65 ubiquitin levels were detected in *VPS35* D620N mouse cortical neurons after 5 hours of AO stimulation. Whole cell lysates were enriched for ubiquitin by incubating with Halo-UBQLN1. Enriched lysates were subjected to immunoblotting with anti-phospho-Ser65 ubiquitin and anti-CISD1 antibodies. MemCode was used as loading control. Whole cell lysates were also subjected to SDS-PAGE and immunoblot analysis with anti-Parkin, anti-phospho-Ser65 Parkin, anti-phospho-Ser111 RAB8A, anti-RAB8A, ant-VPS35 and anti-GAPDH antibodies.

**Figure S3. Ubiquitin Linkage analysis in DIV 16 neurons.**

**A.** Cortical neurons were depolarized with AO (3h) and extracts subject to Ub-AQUA proteomics. Abundance (fmol) or fold increase for individual Ub chain linkage types or pS65-Ub phosphorylation are plotted.

**B.** Same as (A) but extracts were first subjected to sequential Halo-4×UBA^UBQLN1^- and Halo-5xUBA^DSK2^- coupled resin enrichment (see Methods) to isolate ubiquitin and ubiquitylated proteins prior to Ub-AQUA analysis. Error bars represent SEM, n=4. n.d., not determined.

**Figure S4. Proteomic and Biochemical analysis of mouse neurons.**

**A.** Schemata of diGly affinity capture in C57BL/6J primary cortical neurons stimulated with AO for 5 h.

**B.** 5 replicates of E16.5 derived C57Bl/6J primary cortical neurons were stimulated with 10 μM Antimycin A and 1 μM Oligomycin (AO) for 5 h. DMSO as vehicle. Phospho-Ser65 ubiquitin was detected in membrane enriched lysates. GAPDH was used as a loading control.

**C.** Total protein abundance in neurons after treatment with AO. C56BL/6J primary cortical neurons were depolarized with AO (5 h) and membrane enriched lysates were subjected to quantitative proteomics. Fold increase for individual protein is shown in the Volcano plot. The x-axis specifies the fold-changes (FC) and the y-axis specifies the negative logarithm to the base 10 of the t-test p-values (Welch’s t-test (S0=2), corrected for multiple comparison by permutation-based FDR (1%)). Proteins associated with mitochondria (MitoCarta 3.0) or Mitochondrial outer membrane localization are indicated.

**Figure S5. diGLY analysis of neurons stimulated with mitochondrial depolarisation**

**A.** Ranking analysis of least abundant mito Ub sites – MS1- and TMT-based intensity of all diGLY peptides was extracted and the bottom 25 diGLY sites with the smallest relative abundance change upon 5h depolarization is indicated. Error bars represent SEM (n = 5).

**B.** Diagram showing the sites of ubiquitylation in Mitocarta 3.0 and non Mitocarta 3.0 in mouse cortical neurons. Residue numbers for diGLY modified Lys residues are shown.

**Figure S6. Validation of PINK1 activation in AO stimulated Pink1 knockout and wild-type cortical neurons**

**A.** Schematic representation of experimental setup in *Pink1* knockout (KO) and wild-type primary cortical neurons. 3 biological replicates of E 16.5 mouse cortical neurons mouse were cultured for 21 days in vitro. Membrane enrichment performed after mitochondrial depolarisation induced with 10 µM of Antimycin A combined with 1 µM of Oligomycin for 5 h.

**B.** 3 biological replicates of *Pink1* WT and 3 biological replicates of *Pink1* KO primary cortical neurons were stimulated with AO for 5h, DMSO as vehicle. Phospho-Ser65 ubiquitin was detected in membrane enriched lysates. GAPDH was used as a loading control.

**C.** Violin plots for MitoCarta3.0 proteins in wild-type and PINK1^−/−^ primary cortical neurons. Relative protein abundance fold change after 5h depolarization is plotted for the MitoCarta3.0 proteins quantified (black circles). Violin plots represent the distribution and density of the whole dataset (centre line, median; box limits correspond to the first and third quartiles; box whiskers, 1.5x interquartile range; violin limits, minimum and maximum values). Results (*p*-value) of a 2-tailed Mann-Whitney U-test comparing log_2_(AO 5h/UT) between the PINK1^+/+^ and PINK1^-/-^ cell line is indicated.

**D.** Correlation plots (data from Fig 3C&D) for the diGly sites quantified. diGLY-peptide of proteins associated with mitochondria (MitoCarta 3.0) or Mitochondrial outer membrane localization are indicated. Light coloured areas indicate PINK1-dependent or -independent sites.

**E.** Principal component analysis of diGLY proteomics data obtained for the 11-plex experiment described in (A).

**Figure S7. Biochemical analysis of ubiquitylated targets in AO stimulated C56BL/6J cortical neurons.**

Membrane-enriched lysates from C57BL/6J cortical neurons, after 5 hours of AO stimulation, were subjected to ubiquitin capture using TUBE and multiDSK (mDSK) pull-down assay prior to immunoblotting with indicated antibodies. Ubiquitylated forms of CPT1α and CISD1 were detected after mitochondrial depolarisation. Anti-pSer65 Ubiquitin and Ubiquitin antibodies were used as controls. Targets were classified using MitoCarta 3.0

**Figure S8. Analysis of CPT1α in PINK1 KO neurons and SH-SY5Y cells**

**A.** Membrane-enriched lysates of PINK1 WT and KO cortical neurons after 5 h of AO stimulation, were subjected to ubiquitylated-protein capture by Halo-multiDSK (mDSK), prior to immunoblot with anti-CPT1α, anti-CISD1, anti-phosphoSer65 Ubiquitin and anti-Ubiquitin antibodies.

**B.** Timecourse analysis of PINK1-Parkin pathway in SH-SY5Y cell lines. Ubiquitylated forms of CPT1α, CISD1 were found starting from 3 h of AO stimulation in SH-SY5Y cell lines, by using multiDSK ubiquitin pulldown assay. PhosphoSer65 Ubiquitin shows biochemical activation of PINK1.Total ubiquitin is used as loading control. Detection Parkin expression and Parkin Ser65 phosphorylation (Parkin^pS65^) upon AO stimulation (3h, 5h, 9 h, 12h, 18h and 24h). PINK1 stabilisation and OPA1 cleavage are observed after mitochondrial depolarization. GAPDH is used as a loading control.

**Figure S9. Purification of recombinant Parkin substrates and proteins**

**A.** 280 nm UV absorption trace for Cpt1α resolved on a Superdex 200 16/60 size exclusion column (black), with bounds of collected fractions in blue (top panel). Cpt1α fractions under UV absorption peaks resolved on a 4-20 % SDS-PAGE gel and stained using Coomassie brilliant blue (bottom panel). **B.** Purified parkin targets for MBP-CamK2α, MAO-A and GST Miro1; **C.** MBP-CamK2β, MAO-B, Fam213a, SNX3 and ACSL1 (45-end); in both cases 1 μg of the final recombinant protein stock was resolved using 4-20 % SDS-PAGE and stained using Coomassie brilliant blue.

**Figure S10. Ubiquitylation of Parkin substrates**

**A.** Recombinant MAO-A and MAO-B proteins, **B.** FAM213A and **C.** ACSL1 were assessed for ubiquitylation by Parkin. For ubiquitylation assays Parkin was first activated using a 30 min kinase reaction at 37 °C in the presence or absence of wild type (WT) or kinase inactive (KI) TcPink1. Ubiquitylation reactions were started by the addition of the target protein, Flag-Ubiquitin, His-UbE1 and UbE2L3. Ubiquitylation reactions were stopped after a 30 minutes incubation at 37 °C unless stated otherwise by the addition of 4 x LDS loading buffer. Target proteins were resolved on a 4 to 12 % Bis-Tris gels, transferred to nitrocellulose membranes, and immunoblotted using anti-Flag, anti-His, and anti-targeted protein respectively. Membranes were imaged using LiCor.

**Figure S11. PINK1 and Parkin independent ubiquitylation sites**

**A.** Ranking analysis of most abundant Ub sites not associated with MitoCarta3.0 (data from Fig.2A) – MS1- and TMT-based intensity of all diGLY peptides was extracted and the top 50 diGLY sites with the largest relative abundance change upon 5h depolarization is indicated. Error bars represent SEM (n = 5).

**B.** Recombinant SNX3 protein was assessed for ubiquitylation by Parkin and relative mass spectrometry analysis of ubiquitylation sites in wild-type and PINK1 knockout neurons. Kgg fold change to untreated wild-type cells (data from FigS6A) is indicated. Error bars represent SEM, n = 3, 3, 3, 2.

**C.** Recombinant CAMK2 α protein was assessed for ubiquitylation by Parkin and relative mass spectrometry analysis of ubiquitylation sites in wild-type and PINK1 knockout neurons. Kgg fold change to untreated wild-type cells (data from FigS6A) is indicated. Error bars represent SEM, n = 3, 3, 3, 2.

**D.** Recombinant CAMK2 β protein was assessed for ubiquitylation by Parkin \and relative mass spectrometry analysis of ubiquitylation sites in wild-type and PINK1 knockout neurons. Kgg fold change to untreated wild-type cells (data from FigS6A) is indicated. Error bars represent SEM, n = 3, 3, 3, 2.

**Figure S12. Motif analysis for Parkin substrates.**

**A.** Multiple sequence alignment analysis of human sequences (−7 residues to +7 residues of Lys) across common Parkin substrates identified. CISD1 Lys104 and TOMM70 Lys604 were excluded from analysis since Lys residues were located within 5 amino acids of C-terminus.

**B.** Sequence Logo Analysis showed no putative targeting sequence for Parkin-directed Lysine ubiquitylation. Performed using PhosphoSitePlus v6.5.9.3 (https://www.phosphosite.org/sequenceLogoAction.action) online tool.

**Figure S13. DJ1 sites regulated by PINK1**

**A.** Multiple sequence alignment across species encompassing the ubiquitylation sites of DJ1 identified by mass spectrometry. in the figure.

**B.** Kgg fold change to untreated wild-type cells (data from FigS6A) is indicated for PARK7 lysine 93 ubiquitylation. Error bars represent SEM, n = 3, 3, 3, 2.

## ACKNOWLEDGEMENTS

This work was supported by a Wellcome Trust Senior Research Fellowship in Clinical Science (210753/Z/18/Z to M.M.K.M); the Rosetrees Trust (PhD Studentship to O.A.); NIH (RO1 NS083524 to J.W.H.), the Michael J. Fox Foundation (J.W.H. and M.M.K.M.), and a generous gift from Ned Goodnow (J.W.H.). M.L.R. was recipient of a Dundee Research Interest Group (DRIG) studentship, Dundee School of Life Sciences Scholarship, and Bath University Alumni Placement Grant. We thank Joao Paulo for assistance with proteomics (Harvard Medical School). We are grateful to Gail Gilmour and Shauna Channon for mouse genotyping; the Dundee School of Life Science Animal Unit Staff (co-ordinated by Don Tennant and Carol Clacher); the sequencing service (School of Life Sciences, University of Dundee); Axel Knebel for expression and generation of TUBE proteins (MRC PPU); the MRC PPU tissue culture team (co-ordinated by Edwin Allen) and MRC PPU Reagents and Services antibody teams (co-ordinated by James Hastie).

