## Supplemental Figures 1-13 for "Global ubiquitylation analysis of mitochondria in primary neurons identifies physiological Parkin targets following activation of PINK1"

### Figure S1

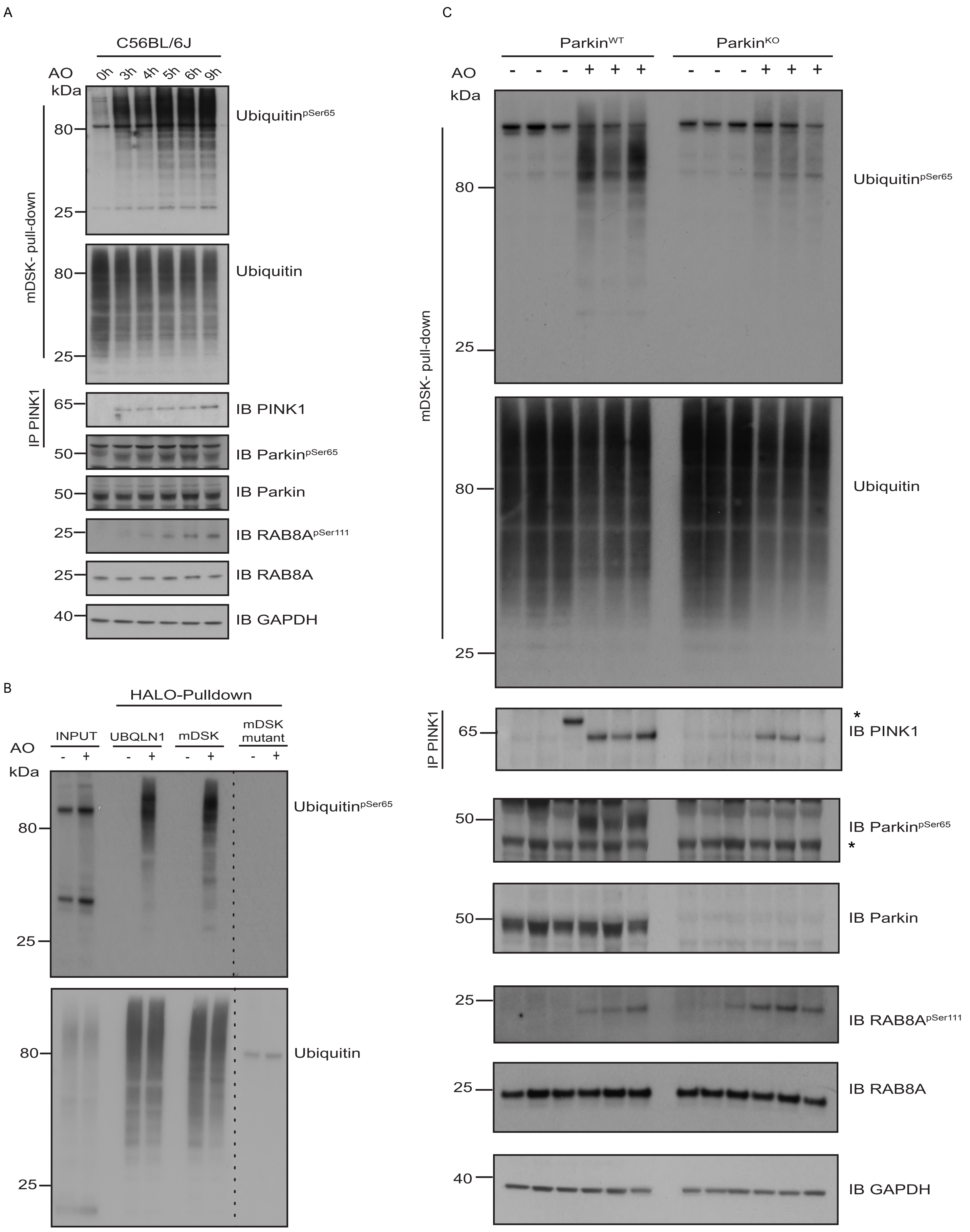

A
C56BL/6J
AO
0h 3h 4h 5h 6h 9h
kDa
80
25
Ubiquitin pSer65
80
25
Ubiquitin
mDSK- pull-down
IP PINK1
65
IB PINK1
50
IB Parkin pSer65
50
IB Parkin
25
IB RAB8A pSer111
25
IB RAB8A
40
IB GAPDH
B
HALO-Pulldown
INPUT
UBQLN1
mDSK
mDSK mutant
AO
- + - + - + - +
kDa
80
25
Ubiquitin pSer65
80
25
Ubiquitin
C
Parkin WT
Parkin KO
AO
- - - + + + - - - + + +
kDa
80
25
Ubiquitin pSer65
80
25
Ubiquitin
mDSK- pull-down
IP PINK1
65
IB PINK1
*
50
IB Parkin pSer65
*
50
IB Parkin
25
IB RAB8A pSer111
25
IB RAB8A
40
IB GAPDH

### Figure S2

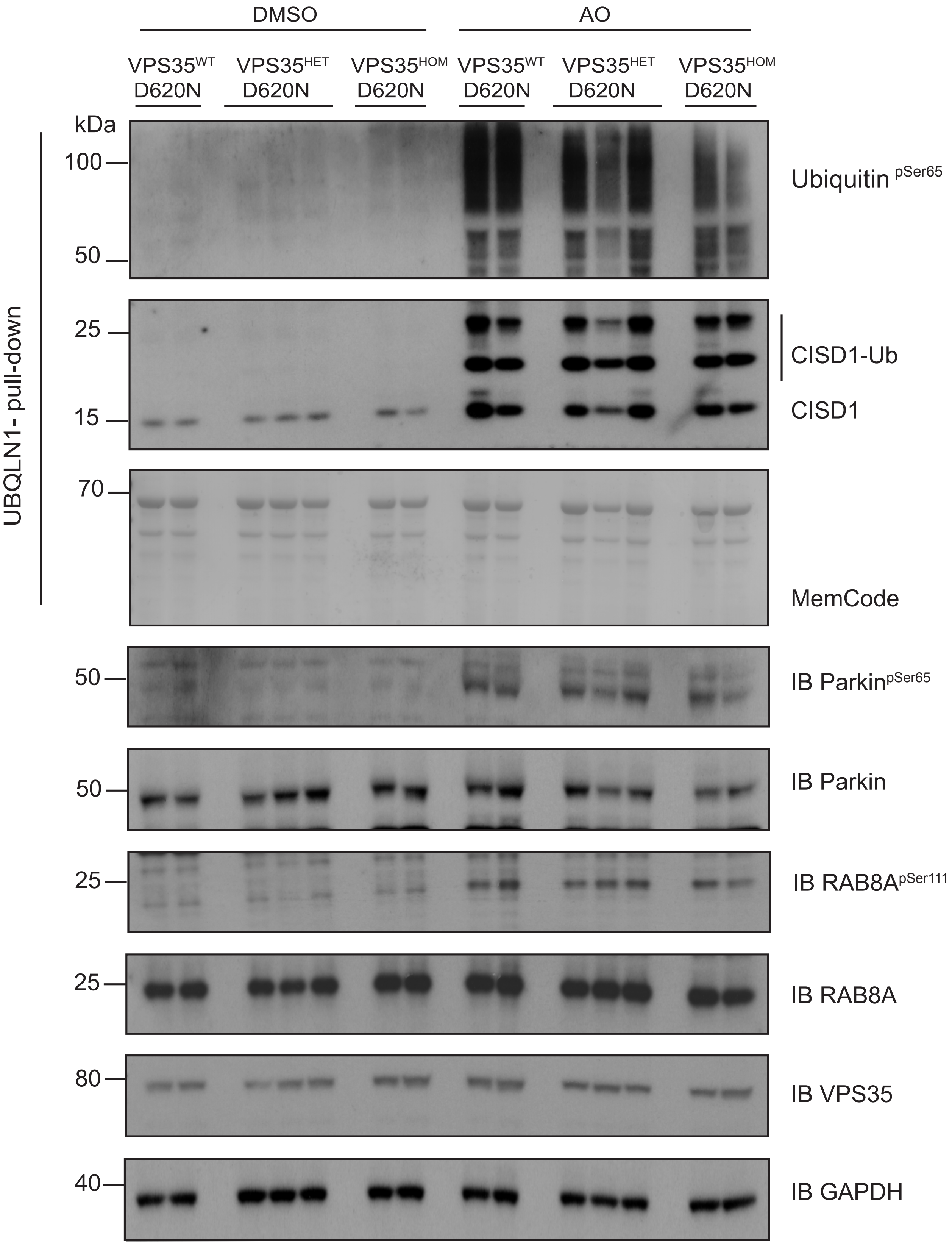
DMSO
AO
VPS35WT D620N
VPS35HET D620N
VPS35HOM D620N
VPS35WT D620N
VPS35HET D620N
VPS35HOM D620N
kDa
100
50
25
15
70
50
50
25
25
80
40
UBQLN1- pull-down
Ubiquitin pSer65
CISD1-Ub
CISD1
MemCode
IB ParkinpSer65
IB Parkin
IB RAB8ApSer111
IB RAB8A
IB VPS35
IB GAPDH

### Figure S3

A

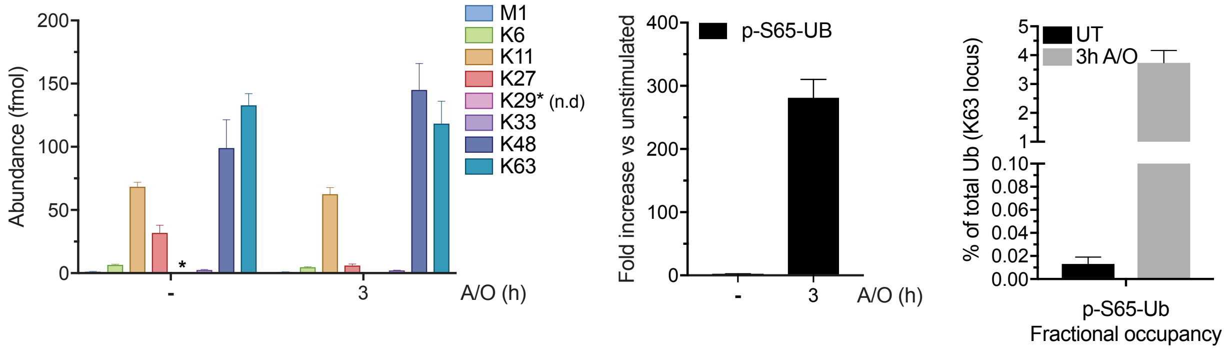
Abundance (fmol)
M1
K6
K11
K27
K29* (n.d)
K33
K48
K63
A/O (h)
p-S65-UB
Fold increase vs unstimulated
UT
3h A/O
% of total Ub (K63 locus)
p-S65-Ub
Fractional occupancy

B

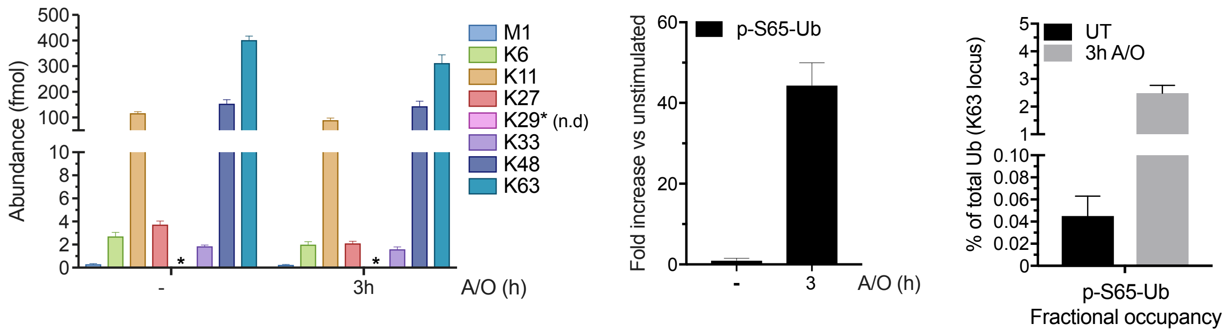
Abundance (fmol)
M1
K6
K11
K27
K29* (n.d)
K33
K48
K63
A/O (h)
p-S65-Ub
Fold increase vs unstimulated
UT
3h A/O
% of total Ub (K63 locus)
p-S65-Ub
Fractional occupancy

### Figure S4

A

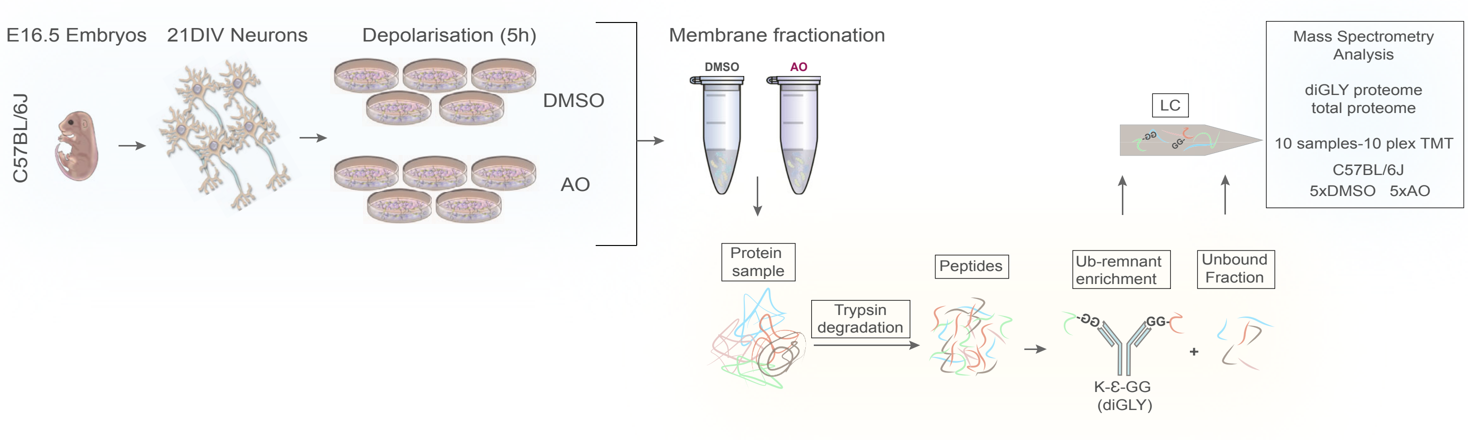


B

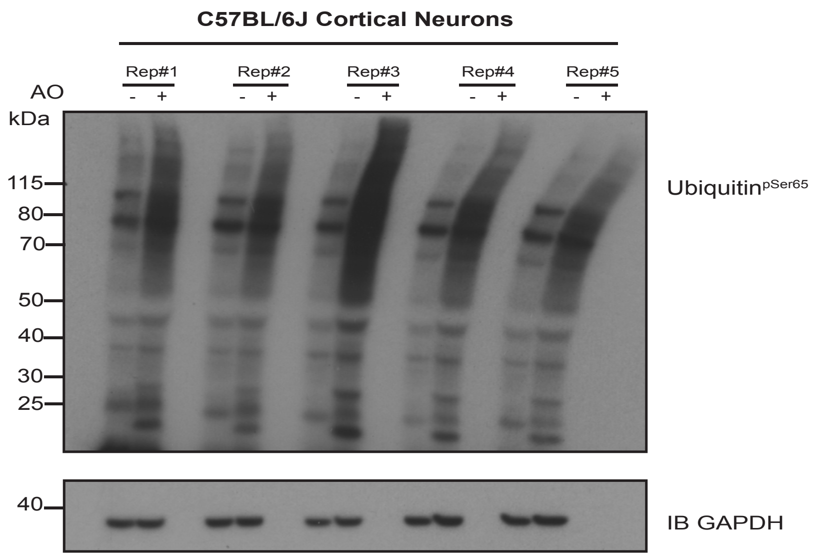


C

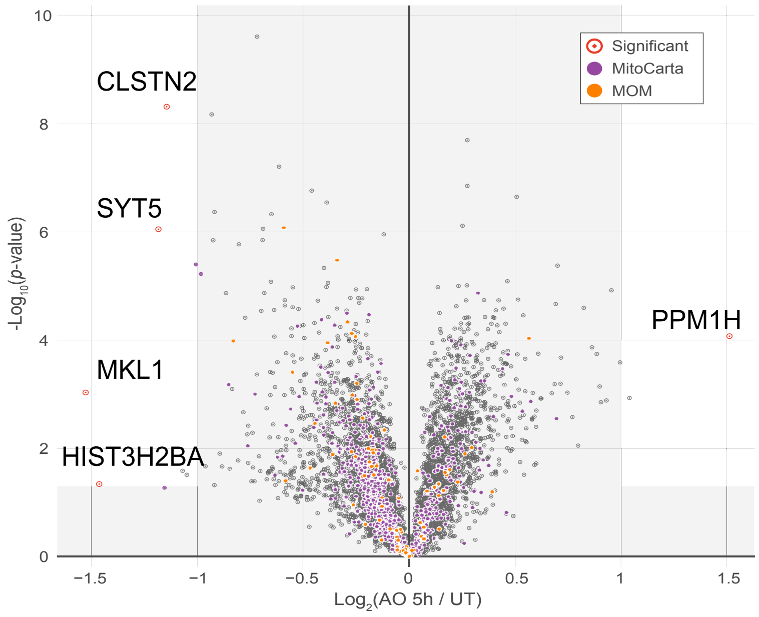

### Figure S5

A

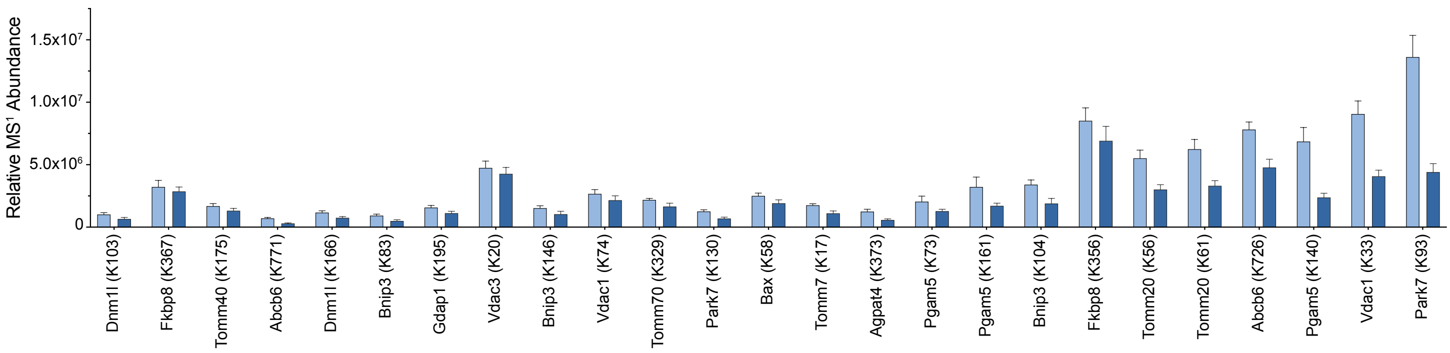

Relative MS¹ Abundance
1.5x10⁷
1.0x10⁷
5.0x10⁶
0
Dnm1l (K103)
Fkbp8 (K367)
Tomm40 (K175)
Abcb6 (K771)
Dnm1l (K166)
Bnip3 (K83)
Gdap1 (K195)
Vdac3 (K20)
Bnip3 (K146)
Vdac1 (K74)
Tomm70 (K329)
Park7 (K130)
Bax (K58)
Tomm7 (K17)
Agpat4 (K373)
Pgam5 (K73)
Pgam5 (K161)
Bnip3 (K104)
Fkbp8 (K356)
Tomm20 (K56)
Tomm20 (K61)
Abcb6 (K726)
Pgam5 (K140)
Vdac1 (K33)
Park7 (K93)


B

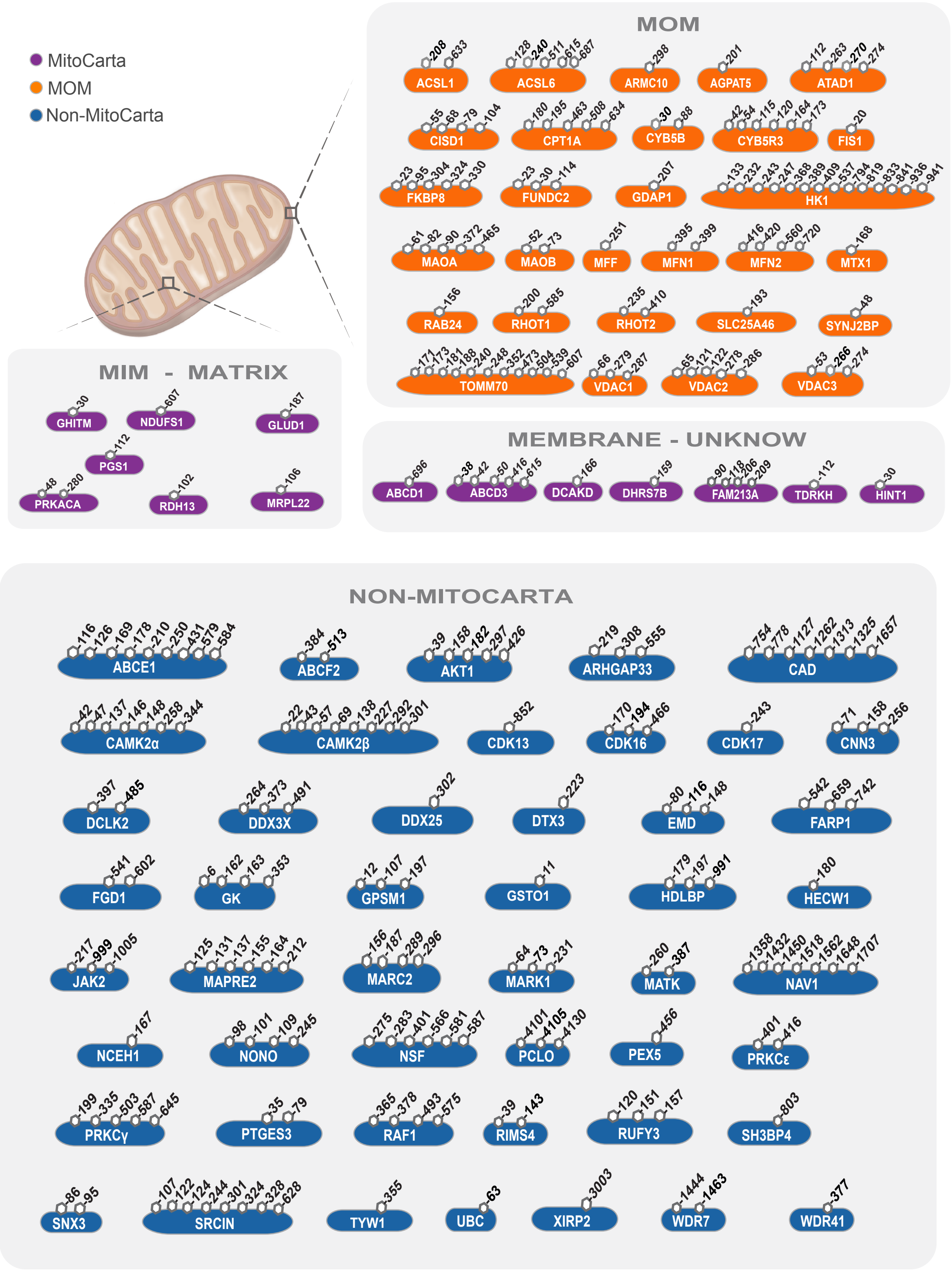

MitoCarta
MOM
Non-MitoCarta
MOM
ACSL1
ACSL6
ARMC10
AGPAT5
ATAD1
CISD1
CPT1A
CYB5B
CYB5R3
FIS1
FKBP8
FUNDC2
GDAP1
HK1
MAOA
MAOB
MFF
MFN1
MFN2
MTX1
RAB24
RHOT1
RHOT2
SLC25A46
SYNJ2BP
TOMM70
VDAC1
VDAC2
VDAC3
MIM - MATRIX
GHITM
NDUFS1
GLUD1
PGS1
PRKACA
RDH13
MRPL22
MEMBRANE - UNKNOW
ABCD1
ABCD3
DCAKD
DHRS7B
FAM213A
TDRKH
HINT1
NON-MITOCARTA
ABCE1
ABCF2
AKT1
ARHGAP33
CAD
CAMK2α
CAMK2β
CDK13
CDK16
CDK17
CNN3
DCLK2
DDX3X
DDX25
DTX3
EMD
FARP1
FGD1
GK
GPSM1
GSTO1
HDLBP
HECW1
JAK2
MAPRE2
MARC2
MARK1
MATK
NAV1
NCEH1
NONO
NSF
PCLO
PEX5
PRKCε
PRKCγ
PTGES3
RAF1
RIMS4
RUFY3
SH3BP4
SNX3
SRCIN
TYW1
UBC
XIRP2
WDR7
WDR41

### Figure S6

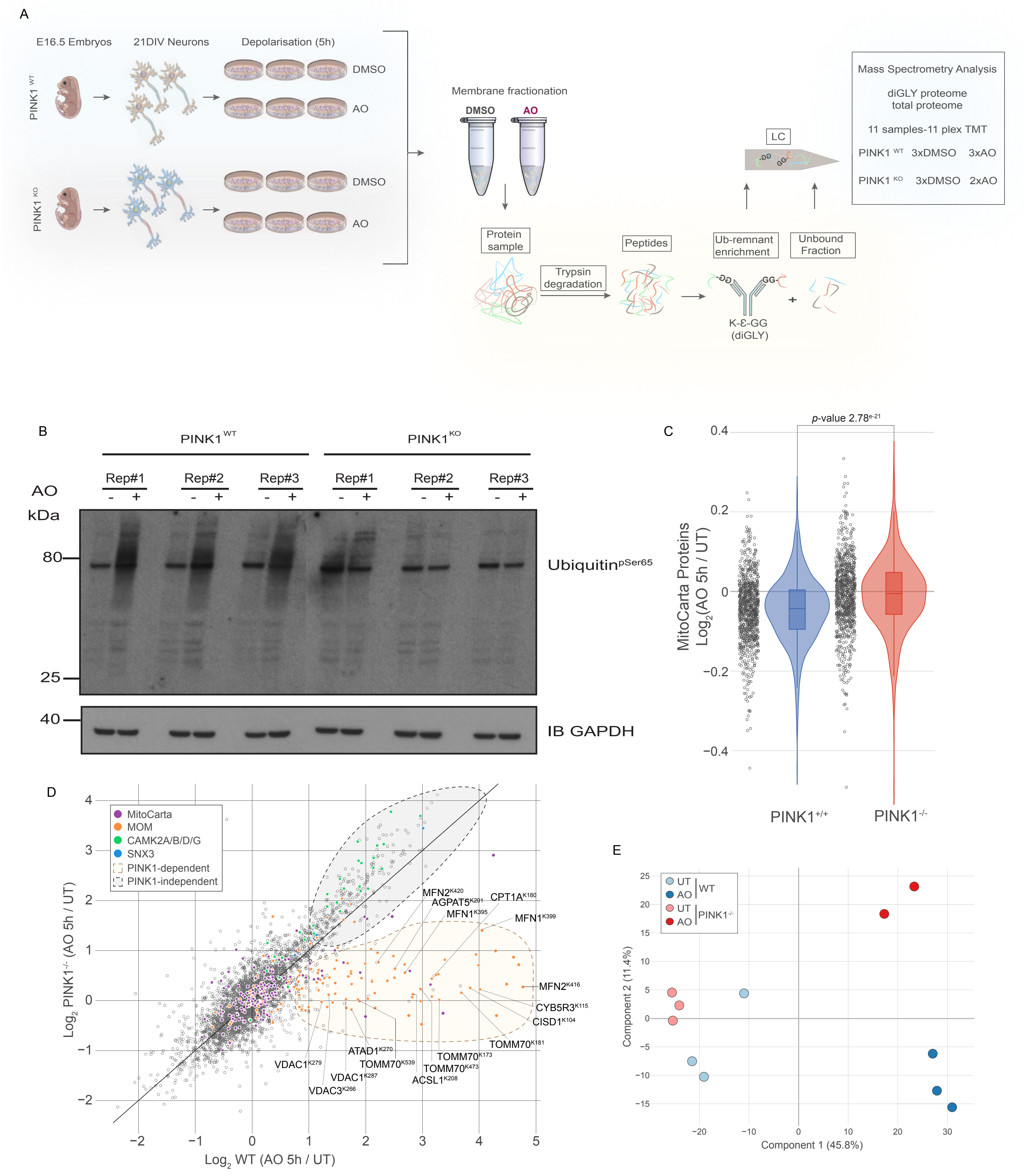
A
E16.5 Embryos
21DIV Neurons
Depolarisation (5h)
PINK1 WT
PINK1 KO
DMSO
AO
DMSO
AO
Membrane fractionation
DMSO
AO
Protein sample
Trypsin degradation
Peptides
Ub-remnant enrichment
Unbound Fraction
K-Ɛ-GG (diGLY)
LC
Mass Spectrometry Analysis
diGLY proteome
total proteome
11 samples-11 plex TMT
PINK1 WT 3xDMSO 3xAO
PINK1 KO 3xDMSO 2xAO
B
PINK1 WT
PINK1 KO
Rep#1
Rep#2
Rep#3
Rep#1
Rep#2
Rep#3
AO
kDa
80
25
40
Ubiquitin pSer65
IB GAPDH
C
p-value 2.78e-21
MitoCarta Proteins Log2(AO 5h / UT)
0.4
0.2
0
−0.2
−0.4
PINK1+/+
PINK1-/-
D
MitoCarta
MOM
CAMK2A/B/D/G
SNX3
PINK1-dependent
PINK1-independent
Log2 PINK1-/- (AO 5h / UT)
Log2 WT (AO 5h / UT)
MFN2 K420
AGPAT5 K201
CPT1A K180
MFN1 K395
MFN1 K399
MFN2 K416
CYB5R3 K115
CISD1 K104
TOMM70 K181
ATAD1 K270
TOMM70 K173
VDAC1 K279
TOMM70 K539
TOMM70 K473
VDAC1 K287
ACSL1 K208
VDAC3 K266
E
UT
AO
WT
UT
AO
PINK1-/-
Component 2 (11.4%)
Component 1 (45.8%)

### Figure S7

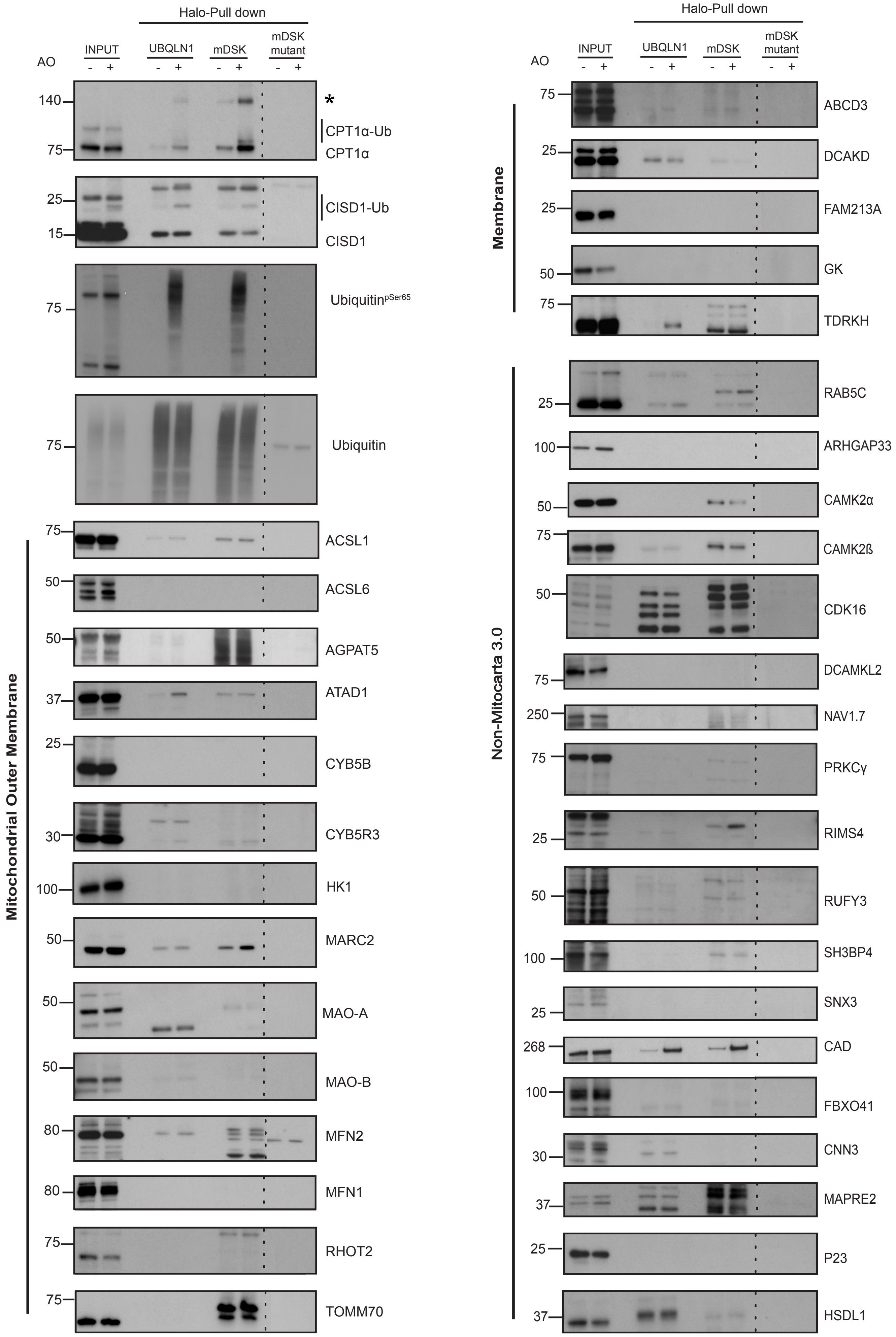

Halo-Pull down
INPUT
UBQLN1
mDSK
mDSK mutant
AO
- +
*
CPT1α-Ub
CPT1α
CISD1-Ub
CISD1
Ubiquitin$^{pSer65}$
Ubiquitin
Mitochondrial Outer Membrane
ACSL1
ACSL6
AGPAT5
ATAD1
CYB5B
CYB5R3
HK1
MARC2
MAO-A
MAO-B
MFN2
MFN1
RHOT2
TOMM70
Membrane
ABCD3
DCAKD
FAM213A
GK
TDRKH
Non-Mitocarta 3.0
RAB5C
ARHGAP33
CAMK2α
CAMK2ß
CDK16
DCAMKL2
NAV1.7
PRKCγ
RIMS4
RUFY3
SH3BP4
SNX3
CAD
FBXO41
CNN3
MAPRE2
P23
HSDL1

### Figure S8

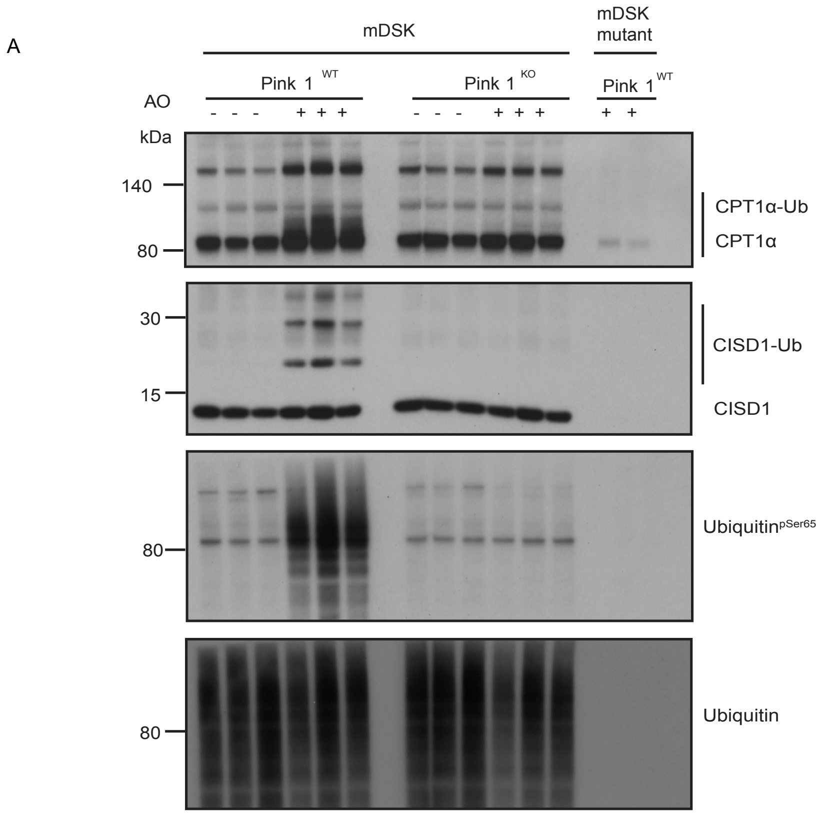
A
mDSK
mDSK mutant
Pink 1 WT
Pink 1 KO
Pink 1 WT
AO
- - - + + + - - - + + + + +
kDa
140
80
CPT1α-Ub
CPT1α
30
15
CISD1-Ub
CISD1
80
UbiquitinpSer65
80
Ubiquitin

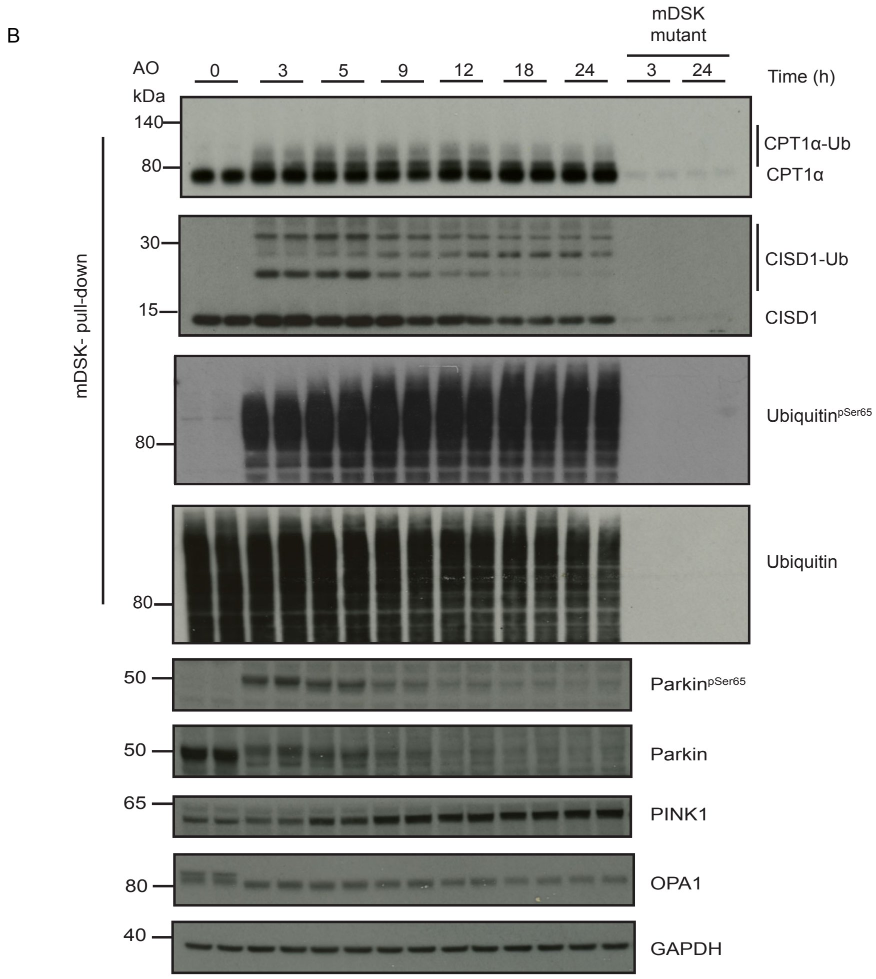
B
mDSK mutant
AO
0 3 5 9 12 18 24 3 24
Time (h)
kDa
140
80
CPT1α-Ub
CPT1α
30
15
CISD1-Ub
CISD1
80
UbiquitinpSer65
80
Ubiquitin
mDSK- pull-down
50
ParkinpSer65
50
Parkin
65
PINK1
80
OPA1
40
GAPDH

### Figure S9

A

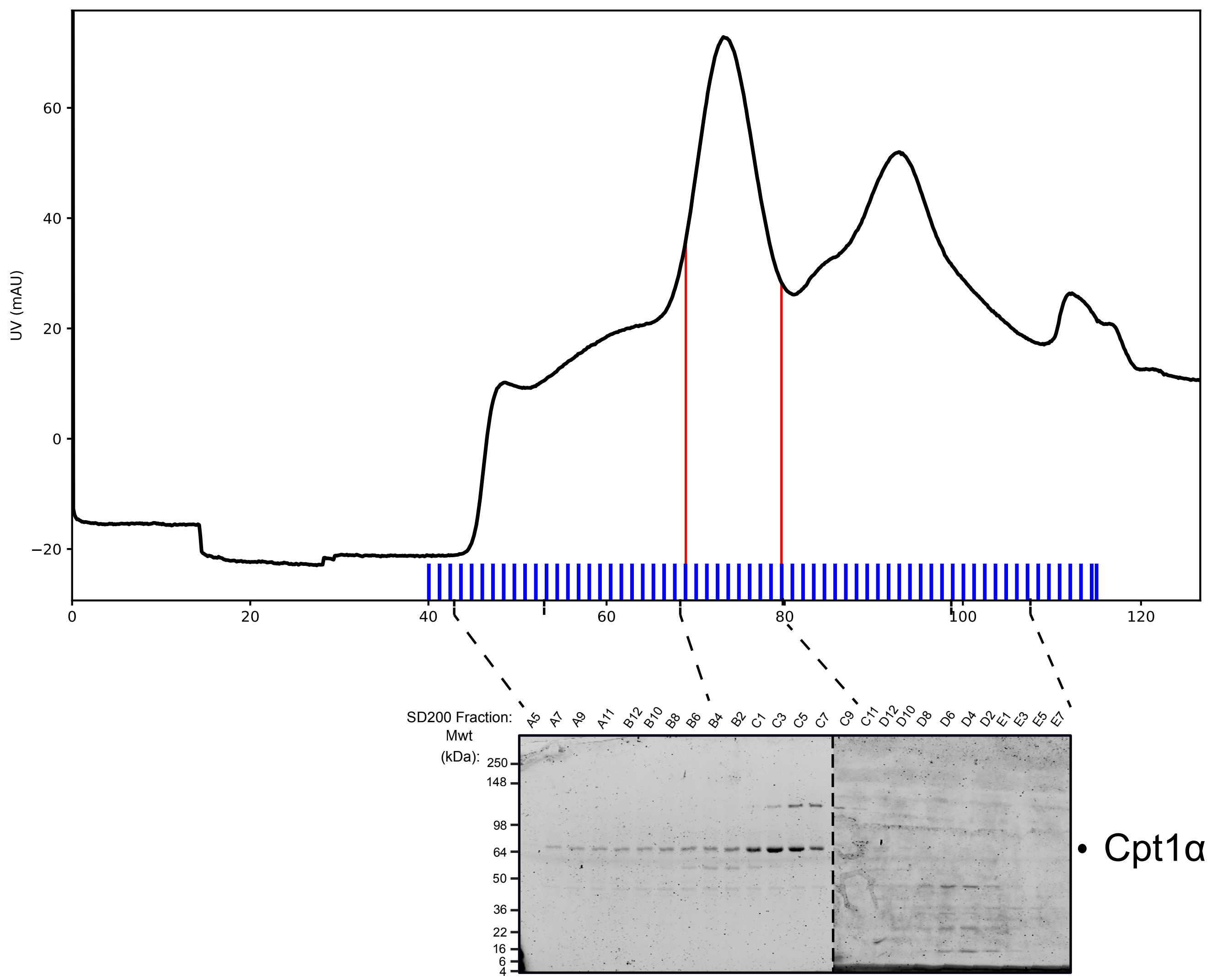


B

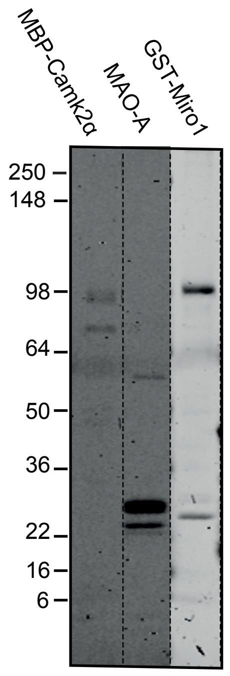


C

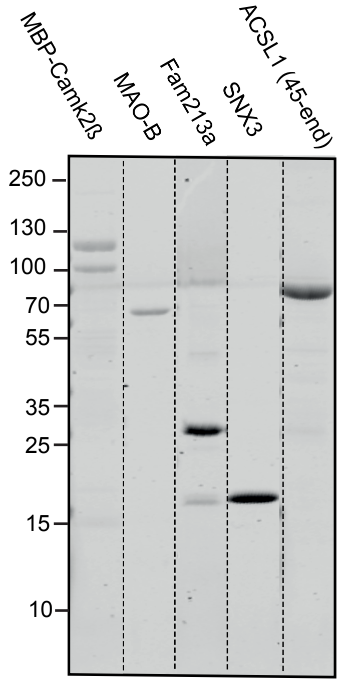

### Figure S10

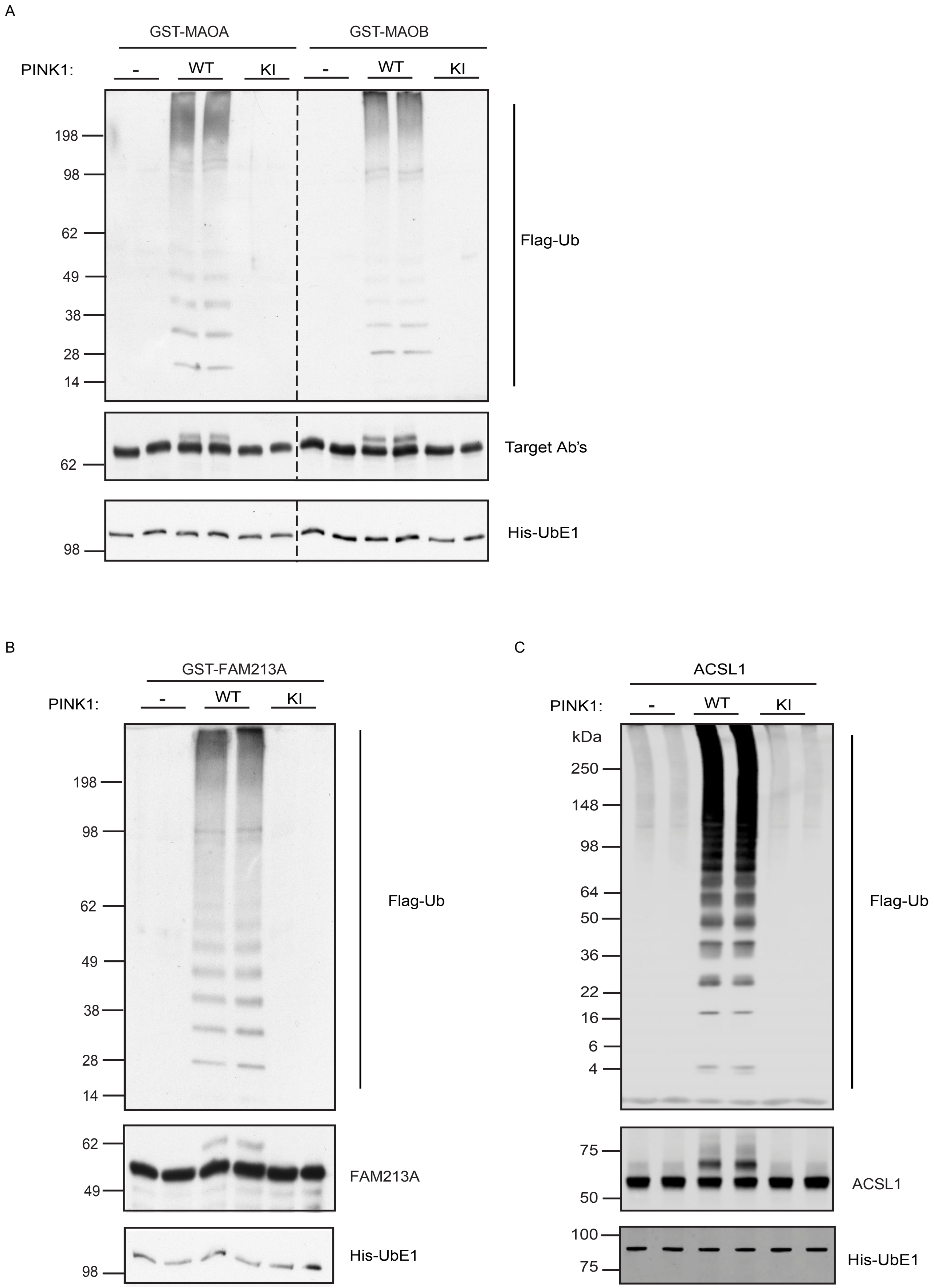
A
GST-MAOA
GST-MAOB
PINK1:
-
WT
KI
198
98
62
49
38
28
14
Flag-Ub
Target Ab's
His-UbE1
B
GST-FAM213A
FAM213A
C
ACSL1
kDa
250
148
64
50
36
22
16
6
4
75
100

### FigureS11

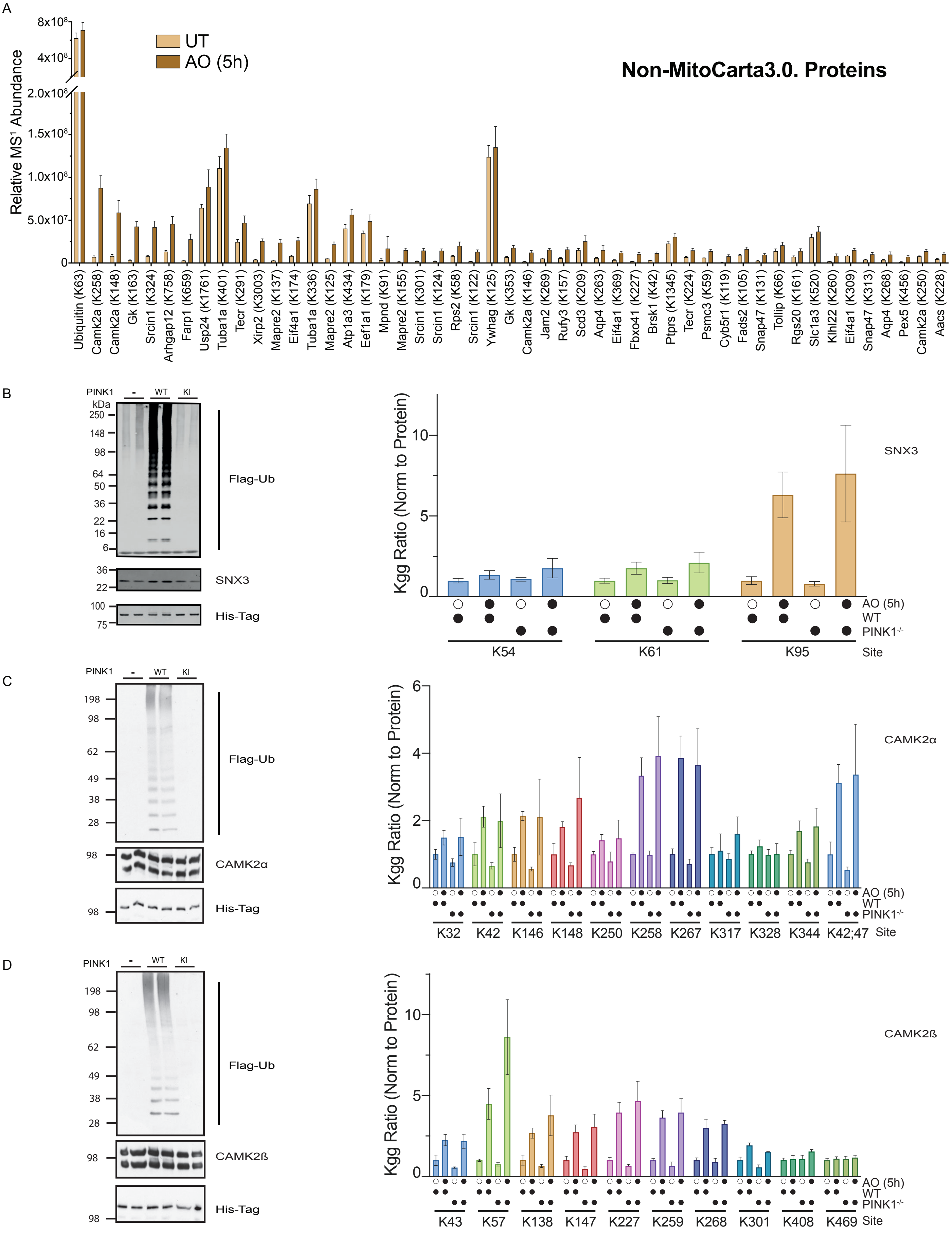
A
UT
AO (5h)
Non-MitoCarta3.0. Proteins
Relative MS1 Abundance
8x10^8
4x10^8
2.0x10^8
1.5x10^8
1.0x10^8
5.0x10^7
0
Ubiquitin (K63)
Camk2a (K258)
Camk2a (K148)
Gk (K163)
Srcin1 (K324)
Arhgap12 (K758)
Farp1 (K659)
Usp24 (K1761)
Tuba1a (K401)
Tecr (K291)
Xirp2 (K3003)
Mapre2 (K137)
Eif4a1 (K174)
Tuba1a (K336)
Mapre2 (K125)
Atp1a3 (K434)
Eef1a1 (K179)
Mpnd (K91)
Mapre2 (K155)
Srcin1 (K301)
Srcin1 (K124)
Rps2 (K58)
Srcin1 (K122)
Ywhag (K125)
Gk (K353)
Camk2a (K146)
Jam2 (K269)
Rufy3 (K157)
Scd3 (K209)
Aqp4 (K263)
Eif4a1 (K369)
Fbxo41 (K227)
Brsk1 (K42)
Ptprs (K1345)
Tecr (K224)
Psmc3 (K59)
Cyb5r1 (K119)
Fads2 (K105)
Snap47 (K131)
Tollip (K66)
Rgs20 (K161)
Slc1a3 (K520)
Klhl22 (K260)
Eif4a1 (K309)
Snap47 (K313)
Aqp4 (K268)
Pex5 (K456)
Camk2a (K250)
Aacs (K228)
B
PINK1
-
WT
KI
kDa
250
148
98
64
50
36
22
16
6
Flag-Ub
36
22
SNX3
100
75
His-Tag
Kgg Ratio (Norm to Protein)
10
5
0
SNX3
AO (5h)
WT
PINK1-/-
K54
K61
K95
Site
C
PINK1
-
WT
KI
198
98
62
49
38
28
Flag-Ub
98
CAMK2α
98
His-Tag
Kgg Ratio (Norm to Protein)
6
4
2
0
CAMK2α
AO (5h)
WT
PINK1-/-
K32
K42
K146
K148
K250
K258
K267
K317
K328
K344
K42;47
Site
D
PINK1
-
WT
KI
198
98
62
49
38
28
Flag-Ub
98
CAMK2ß
98
His-Tag
Kgg Ratio (Norm to Protein)
10
5
0
CAMK2ß
AO (5h)
WT
PINK1-/-
K43
K57
K138
K147
K227
K259
K268
K301
K408
K469
Site

### Figure S12

A

| | |
|---|---|
| ACSL1 207 | VDKPEKA**K**LLLEGVE |
| AGPAT5 201 | QRGLAVL**K**HVLTPRI |
| ATAD1 112 | DTVILPI**K**KKHLFEN |
| ATAD1 270 | KQREAIL**K**LILKNEN |
| CPT1 180 | RLPVPAV**K**DTVNRYL |
| CYB5R3 42 | TLESPDI**K**YPLRLID |
| CYB5R3 115 | LVIKVYF**K**DTHPKFP |
| CYB5R3 120 | YFKDTHP**K**FPAGGKM |
| CYB5R3 154 | GLLVYQG**K**GKFAIRP |
| CYB5R3 164 | GLLVYQG**K**GKFAIRP |
| CYB5R3 173 | NPIIRTV**K**SVGMIAG |
| FAM213A 129 | KDFQPYF**K**GEIFLDE |
| FIS1 20 | DLLKFEK**K**FQSEKAA |
| GDAP1 207 | DNVKYLK**K**ILDELEK |
| GHTM 29 | TKASPVV**K**NSITKNQ |
| HK1 77 | SIPDGSE**K**GDFIALD |
| HK1 176 | ITWTKRF**K**ASGVEGA |
| HK1 187 | VEGADVV**K**LLNKAIK |
| HK1 191 | DVVKLLN**K**AIKKRGY |
| HK1 333 | PELLTRG**K**FNTSDVS |
| HK1 738 | EYSLNAG**K**QRYEKMI |
| HK1 785 | TRGIFET**K**FLSQIES |
| HK1 885 | TVKELSP**K**CNVSFLL |
| MFN1 395 | NLLTLDV**K**KKIKEVT |
| MFN1 399 | LDVKKKI**K**EVTEEVA |
| MFN2 416 | ELLAQDY**K**LRIKQIT |
| MFN2 420 | QDYKLRI**K**QITEEVE |
| MFN2 720 | EIAAMNK**K**IEVLDSL |
| RAB24 156 | QLFETSS**K**TGQSVDE |
| RHOT1 187 | EMKPACI**K**ALTRIFK |
| RHOT2 409 | EKRLDQE**K**GQTQRSV |
| SYNJ2BP 48 | GIYVSRI**K**ENGAAAL |
| TDRKH 112 | NEIGAIE**K**AVIWPQY |
| TOMM70 170 | FEQLQKW**K**EVAQDCT |
| TOMM70 178 | EVAQDCT**K**AVELNPK |
| TOMM70 185 | KAVELNP**K**YVKALFR |
| TOMM70 245 | EKAKEKY**K**NREPLMP |
| TOMM70 470 | KGFEEVI**K**KFPRCAE |
| TOMM70 536 | RGLELIS**K**AIEIDNK |
| VDAC1 53 | SANTETT**K**VTGSLET |
| VDAC1 266 | LSALLDG**K**NVNAGGH |
| VDAC1 274 | NVNAGGH**K**LGLGLEF |
| VDAC2 64 | SSNTDTG**K**VTGTLET |
| VDAC2 285 | SINAGGH**K**VGLALEL |
| VDAC3 53 | HAYTDTG**K**ASGNLET |
| VDAC3 266 | LSALIDG**K**NFSAGGH |
| VDAC3 274 | NFSAGGH**K**VGLGFEL |

B

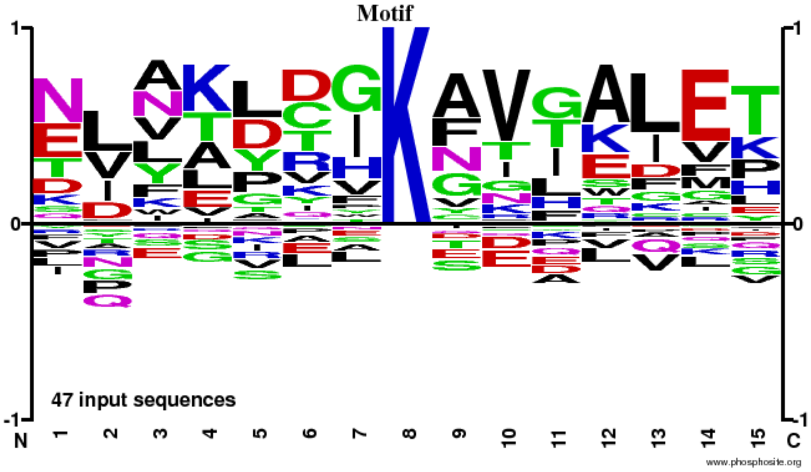

### Figure S13

A

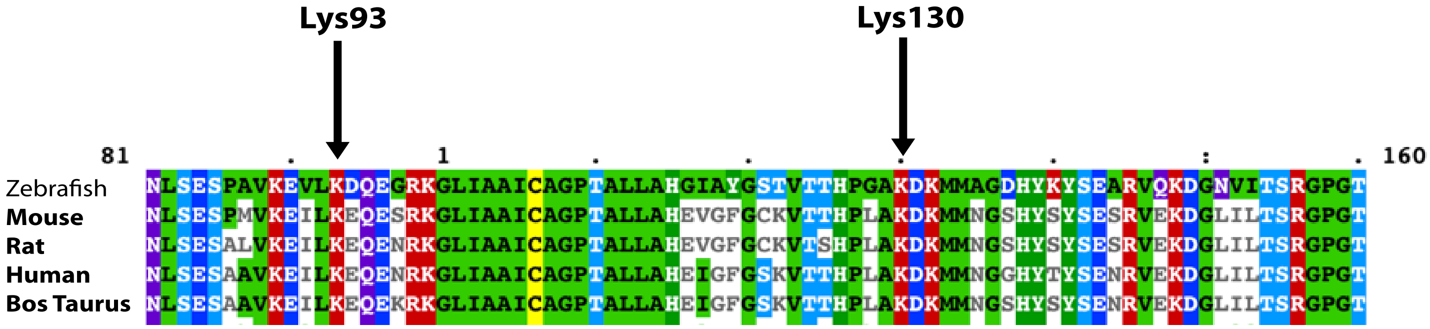


B

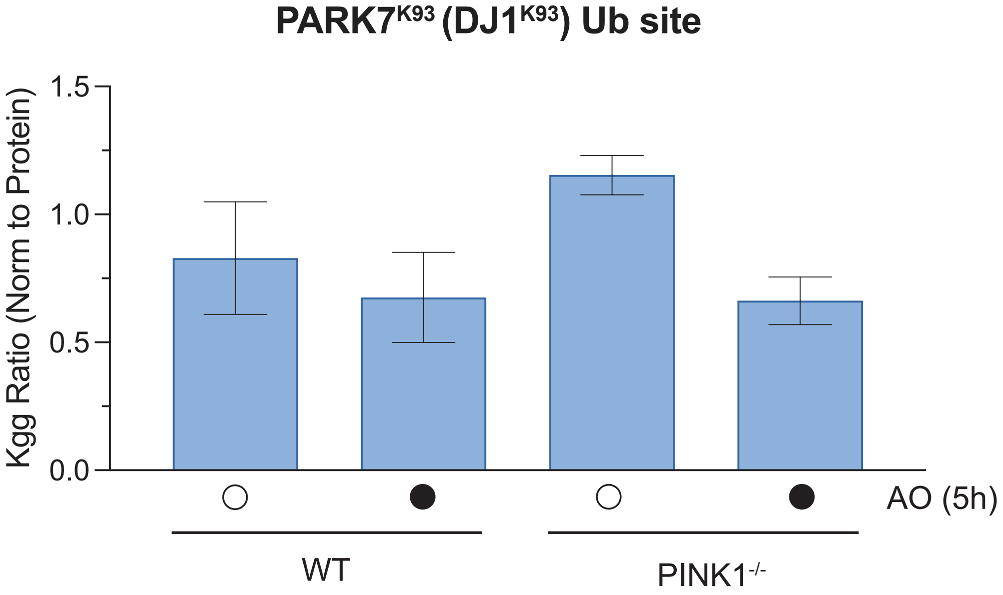
